# A new long-read dog assembly uncovers thousands of exons and functional elements missing in the previous reference

**DOI:** 10.1101/2020.07.02.185108

**Authors:** Chao Wang, Ola Wallerman, Maja-Louise Arendt, Elisabeth Sundström, Åsa Karlsson, Jessika Nordin, Suvi Mäkeläinen, Gerli Rosengren Pielberg, Jeanette Hanson, Åsa Ohlsson, Sara Saellström, Henrik Rönnberg, Ingrid Ljungvall, Jens Häggström, Tomas F. Bergström, Åke Hedhammar, Jennifer R. S. Meadows, Kerstin Lindblad-Toh

## Abstract

Here we present a new high-quality canine reference genome with gap number reduced 41-fold, from 23,836 to 585. Analysis of existing and novel data, RNA-seq, miRNA-seq and ATAC-seq, revealed a large proportion of these harboured previously hidden elements, including genes, promoters and miRNAs. Short-read dark regions were detected, and genomic regions completed, including the DLA, TCR and 366 cancer genes. 10x sequencing of 27 dogs uncovered a total of 22.1 million SNPs, Indels and larger structural variants (SVs). 1.4% overlap with protein coding genes and could provide a source of normal or aberrant phenotypic modifications.

## Background

Man and his best friend the dog share the same environment, most of their genes and a large number of diseases. Dogs have lived alongside people for at least 10,000 years, becoming increasingly domesticated[1,2], and have in the past few hundred years been bred into hundreds of individual breeds. As specific traits such as morphology and behavior have been under selection, unwanted traits and diseases have become more common within particular dog breeds due to drift and/or cosegregation of disease risk alleles with the selected traits.

Because of the favorable genetic structure and the disease predisposition in certain breeds, tools have been generated to allow for trait mapping. For many diseases, variants affecting specific genes have been found by genome wide association studies. This knowledge has been used to understand the biology of both canine and human disease, including both inherited and somatic mutations in cancer. The current canine genome reference, CanFam3.1, is based on Sanger sequencing and was first published in 2005[3]. It has been improved by multiple methods since[4], but still contains 23,876 gaps, many of which are found close to the 5’ end of genes. The accumulation of such genomic regions, difficult to read through with previous methods, is at least partially due to the loss of *PRDM9* in dogs[5] leading to regions with very high GC content. Here, we generated a high-quality canine reference assembly using a combination of Pacific Biosciences (PacBio) long-read sequencing, 10x Genomics Chromium Linked-Reads (henceforth called 10x) and Hi-C proximity ligation. To enhance gene and variant annotation we generated additional WGS, ATAC and cDNA sequencing.

## Results and Discussion

We selected a Swedish 12-year-old German shepherd with no history of genetic disorders, Mischka, as the source for our high-quality reference genome assembly (CanFam_GSD1.0; henceforth called GSD1.0; Additional file 1: Fig. S1). We sequenced the genome using 91X coverage PacBio, 94X of 10x and 48X coverage of Hi-C linked reads. Contigs were assembled with FALCON and scaffolded with 10x and Hi-C linked reads into 39 single-scaffold chromosomes (total 2.35Gb) and 2,159 unplaced scaffolds (total 128.5Mb; Fig. 1a). The latter mainly originating from segmental duplications and centromeric repeats Additional file 1: Fig. S2). Compared to CanFam3.1, the contiguity of GSD1.0 has been improved 41-fold reaching a contig N50 of 14.8Mb (Additional file 1: Fig. S3), with only 367 gaps in the chromosome scaffolds (Additional file 2: Tab. S1). The quantity of sequence with extreme GC content (>90%, 50bp-window) has doubled to 1.7Mb (Fig. 1b), leading to a 14% increase in the average length of CpG islands (1,056bp versus 926bp, P=8.4 × 10^−4^, t-test). Filled gaps were found to have either high GC or repeat content (Fig. 1c).

**Figure 1.**
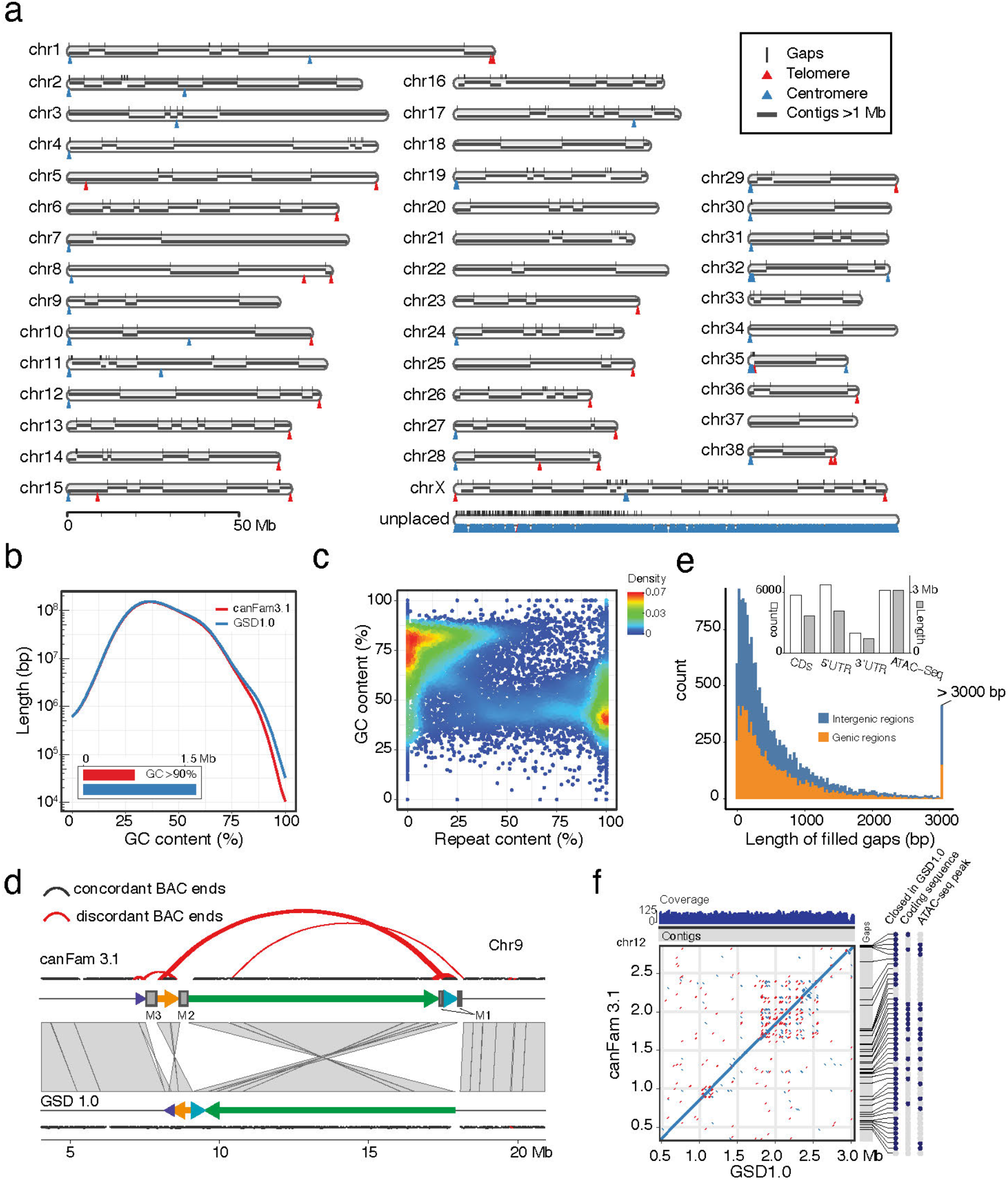
Features of the novel canine assembly. a) GSD1.0 ideogram showing chromosomes, contigs, gaps, centromere and telomere repeats. All unplaced sequences were concatenated into a single scaffold (segmental duplications, 58.1% and centromeric repeats, 30.1%). b) Comparison of GC content (50 bp-window) between GSD1.0 and CanFam3.1. c) Characteristics of the sequences from the gaps closed in GSD1.0. Most of these are either high in GC or repeat content. d) Correction of an inverted region in chromosome 9, which was confirmed by mapping the end sequences of BAC (CH82). 49 discordant pairs (red curves) were found on the edge of rearranged fragments in CanFam3.1. However, these pairs were correctly mapped in GSD1.0 e) Size distribution and overlap with exons and promoters for filled gaps. f) Sequence comparison of DLA on chromosome 12 between CanFam3.1 and GSD1.0.

Using repeatmasker we found 42.7% of the genome to contain repeats, with the three major categories being LINEs (504Mb), SINEs (253Mb) and LTRs (120Mb) (Additional file 1: Fig. S4, Additional file 2: Tab. S2). Based on the position of centromeric repeats, the orientation of chromosomes 27 and 32 was reversed compared to CanFam3.1. Centromeres are now represented at the beginning of the q-arm of 21 autosomes, and 17 autosomes end in telomeric repeats (Fig. 1a). As expected, the X chromosome has telomeric repeats at each end, and a clear centromeric signal at 49.4-49.9Mb. In addition, throughout the genome we found a total of 10 internal centromeric, and 7 internal telomeric repeats. Most of these were also present in the previous genome assemblies and may indicate ancient centromere and telomere positions prior to chromosomal rearrangements. Using HiC and BAC end sequencing data, we identified a sequence belonging on chromosome 18 misassembled on chromosome 9 in CanFam3.1. This sequence is composed of a set of complex structural variants (Fig 1d) and overlaps with the *MAGI2*. Various somatic genetic alterations in this gene are observed in ovarian, breast and colorectal carcinomas[6–8].

Five additional dog genome assemblies have recently been deposited in NCBI (Additional file 2: Tab. S3). We benchmarked GSD1.0 against CanFam3.1 and these assemblies using BUSCO[9] and Iso-Seq cDNA alignments from an unrelated beagle. Compared to CanFam3.1, 25,609 (4.8%) of Iso-Seq reads could be mapped with >5% more bases on GSD1.0 (Additional file 1: Fig. S5), a higher proportion than for the other assemblies (Additional file 2: Tab. S4). GSD1.0 has the second highest BUSCO score for complete genes (95.5%).

A new annotation based on RNA-seq evidence was generated to resolve transcript length and account for the gap closures in GSD1.0. We generated more than 70M nanopore and PacBio full-length cDNA reads from multiple brain and retina regions, and used this in combination with 24 billion public RNA-seq paired reads to annotate 165k transcripts in 29,406 genes. 20,483 genes are potentially protein coding with an open reading frame of >100 amino acids, and 19,586 genes had a significant BLAST hit against proteins in Swissprot or ENSEMBL. We identified 9,356 long noncoding genes (>200 bases and at least 2 exons). The GSD1.0 annotation has a higher number of genes with an ORF fully matching a protein when compared to proteins extracted from CanFam3.1 (Additional file 1: Fig. S6). Gene predictions and non-dog refSeq alignments were used to identify potentially missed genes that did not overlap with our annotation. The result was 874 additional protein coding genes with BLAST evidence, and the BUSCO score for complete genes increased from 98.6% to 99.0%.

We identified 5,743 unique coding exons in the filled CanFam3.1 gaps. 7,468 gaps contained either an exon or promoter sequence as defined by ATAC-seq peaks (Fig 1e). Analysis of these regions revealed eight genes with >80% of their coding sequence in the filled gaps, *PSMA4, CDHR5, SCT, PAOX, UTF1, EFNA2, GPX4,* and *SLC25A22* - which has a role in human cancers (Fig. 2a)[10]. Using a combination of new miRNA-seq reads and public data we identified a conservative set of 719 miRNAs. This included Mirlet-7i, one of the most highly expressed microRNAs in the dataset (Additional file 1: Fig. S7). Mirlet7i has been found to be involved in several diseases, including multiple sclerosis, gastric cancer and breast cancer[11–13].

**Figure 2.**
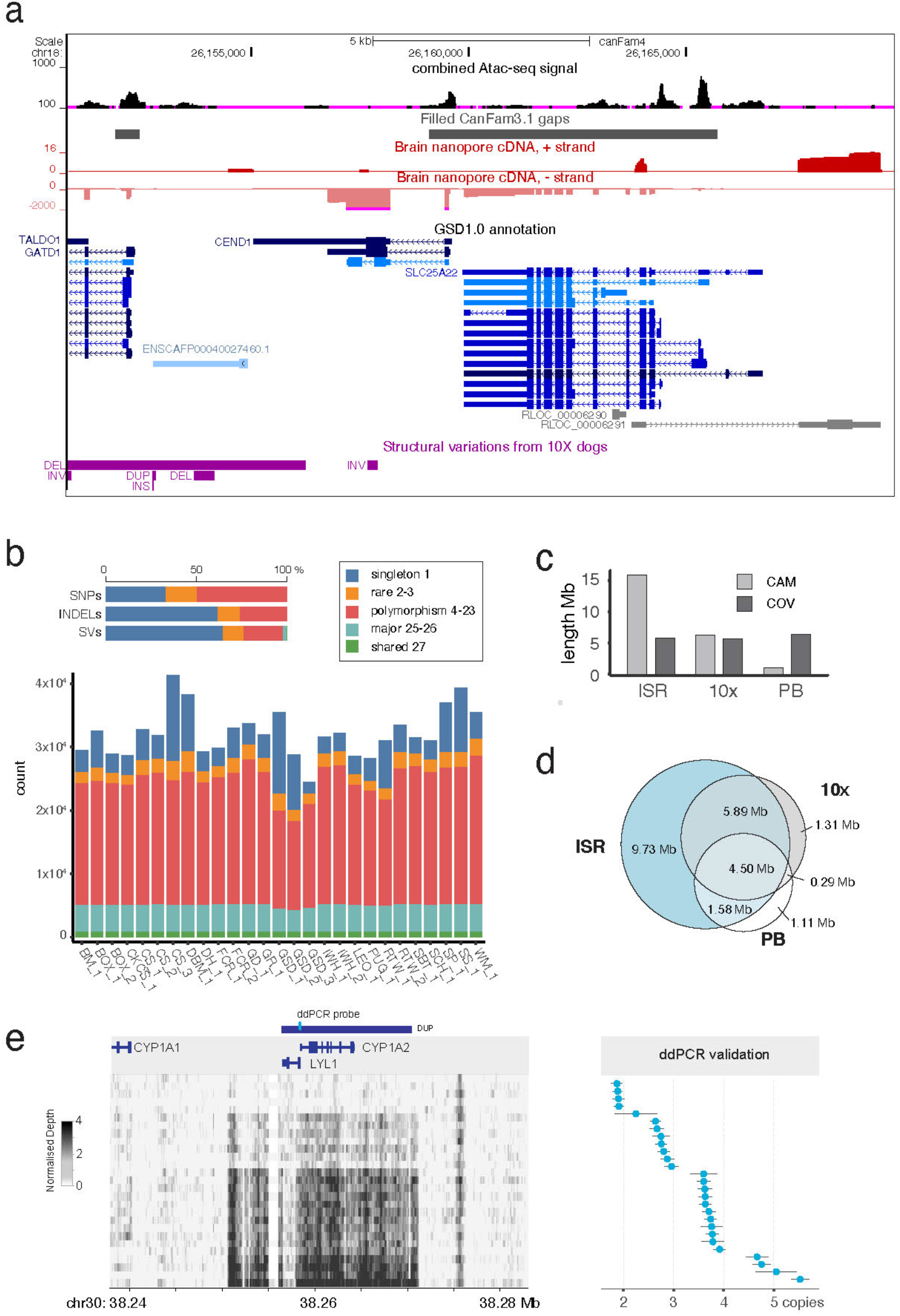
Genome variation, dark regions and genes. a) Representative GSD1.0 annotation from the UCSC track hub highlighting available data and an example of a gene hidden in CanFam3.1. b) SNPs, Indels and structural variations shared among Mischka and the 27 10x sequenced dogs. c) The total length of dark regions detected from Illumina short-reads (ISR), 10x and PacBio sequencing. d) Intersection of dark regions from different datasets. e) Identification and validation of duplication on chromosome 30 containing the *CYP1A2* gene.

In order to improve the dog as a spontaneous model for studying cancer and immunological diseases it is important that key regions are correctly assembled. From COSMIC[14], 282 tier1 and 78 tier2 genes are now completely captured (Additional file 2: Tab. S5), including *HOXD13* and *KLF4*. Based on previous evidence HOXD13 methylation status functions as a prognostic indicator in cancer and deubiquitination of KLF4 promotes breast cancer metastasis (Additional file 1: Fig. S8)[15,16]. Moreover, we verified that regions for Dog Leukocyte Antigen (DLA) positioned on chromosomes 12 (Fig. 1f) and 35 (Additional file 1: Fig. S9a) are contiguous in GSD1.0 (covering 2.58Mb and 0.61Mb, respectively). New coding and potential regulatory sequences previously hidden in gaps in CanFam3.1 were also identified in this region. Furthermore, the T-cell receptor alpha (TRA) and T-cell receptor beta (TRB) loci on chromosome 8 and 16 respectively (Additional file 1: Fig. S9b&c), are both contiguous.

To better understand the genome characteristics and investigate variation across dog breeds, 27 dogs from 19 breeds (Additional file 2: Tab. S6) were selected and sequenced with 10x to a mean depth of 44X (range 30-80X, Fig. 2b). Using Long Ranger we detected 14,953,199 SNPs, 6,958,645 Indels and 217,951 structural variants. Of these 42.1% were only present in one sample, while 57.9% were polymorphic across many individuals. A total of 1.4% (295,112 SNPs and 16,654 SVs) overlapped with protein-coding regions.

With a large number of gaps filled, we set out to study the “dark matter” of the genome[11]: regions that can either not be adequately covered by the sequencing method (COV) or uniquely aligned to the assembly, “camouflaged” (CAM). We defined dark regions in GSD1.0 for Illumina (ISR), 10x, and PacBio (PB) sequencing (see method). COV comprised 5.8Mb, 5.7Mb and 6.4Mb respectively, while CAM comprised 15.9Mb, 6.4Mb and 1.0Mb (Fig. 2c). Intersection showed that while 10x could rescue 11.3Mb dark regions not seen with standard Illumina libraries, significantly more of these regions were covered by PacBio (Fig. 2d). We identified 51,994 short variants in dark regions, including 19,340 intronic and 2,074 exonic variants. Many of these variants were embedded in genes that may be important for morphology or associated with diseases. For example, 14 variants were found within seven intronic *TYRP1* ISR CAM dark regions (Additional file 1: Fig. S10a): a gene linked to brown color in dogs[17]. Likewise, 76 variants were found in *ADCY2* ISR CAM regions (Additional file 1: Fig. S10b). Polymorphisms in this gene have previously been associated with severe chronic obstructive pulmonary disease in humans[18].

To assess the potential role of SVs in gene expression, we selected three segregating deletions and three CNVs for further inspection (Additional file 2: Tab. S7). The largest CNV locus spanned 16.2kb and encompassed potential regulatory regions and two protein coding genes, including *CYP1A2* (Fig 2e). In its homozygous mutant state, a known canine *CYP1A2* premature stop codon (rs852922442; c.1117C>T; R373X)[19,20] results in an array of pharmacokinetic effects, including reduced hepatic drug metabolism[21]. The rs852922442-T variant is present in a range of breeds[22], and was observed in 4/27 10x dogs, but in heterozygous form and not segregating with CNV count. The *CYP1A2* copy number of the 10x individuals was resolved to range from two to five copies (Fig 2e). Differential gene expression analyses using either liver or spleen did not result in any locus wide significant results and so the phenotypic consequence of this expansion has yet to be resolved (CNV=3 vs CNV>3; Additional file 2: Tab. S8). It may be that the effect in this region is subtle, and so not detectable with qPCR, however, *CYP1A2* is an inducible gene and so the true effect may only be observed after a drug challenge[23].

## Conclusion

The GSD1.0 canine genome assembly is highly improved compared to CanFam3.1. We filled ~23,000 gaps, allowing for the completion of a large number of genes, both exons and promoters, as well as for the capture of several additional full-length genes. Key complete regions included the DLA and TCR as well as hundreds of cancer genes. We also analyzed dark regions, generating a catalogue of where the genome might be more challenging to analyse. By combining correct gene models with the presence of both SNPs and SVs, we identify variants we expect can be used to explain large phenotypic differences, disease as well as drug sensitivity in dogs. Our hope is that this will propel the comparison of canine and human genetic disease forward.

## Methods

### Animal samples and WGS for Mischka

A 12-year-old female Swedish German Shepherd with no medical history of genetic disease, Mischka, was selected for the construction of genome assembly. We collected blood for genomic sequencing and tissue samples for Hi-C. Genotyping with the Illumina 170K CanineHD beadchip confirmed Mischka to genetically represent German Shepherd based on the expected inbreeding value (F=0.037, PLINK v1.9), and multidimensional scaling (MDS) analysis with 260 Swedish German Shepherds from a previous study (PLINK v1.9)[24].

High molecular weight (HMW) DNA for PacBio and 10x sequencing was extracted from blood samples using the MagAttract HMW DNA Kit (QIAGEN, USA). The PacBio libraries were prepared using SMRTbell Template Prep Kit 1.0, and then sequenced using 70 SMRT cells on the PacBio Sequel system with v2.1 chemistry. (Fig. S9). We also sequenced HMW DNA of Mischka with Chromium libraries (10x Genomics, USA) on Illumina HiSeq X (2×15bp), to generate a total of 269.75Gb data (~96X).

Three HiC libraries were prepared from Mischka blood samples at Dovetail Genomics (Scotts Valley, Ca, USA) following their standard protocol (https://dovetailgenomics.com/wp-content/uploads/2019/08/Dovetail™-Hi-C-User-Manual_v-1.4-.pdf). Illumina libraries were sequenced on HiSeq X as 2×150bp paired-end reads. We generated a total of 121.47Gb data from three HiC libraries with an estimated sequence coverage of 48.6X (Additional file 2: Tab. S9).

### Assembly construction

PacBio subreads >8Kb were used for *de novo* assembly with FALCON (v0.5.0)[25]. We chose the standard FALCON assembly instead of the “UNZIP” version, since the latter is recommended for samples with a higher heterozygosity. The assembly sequences were polished by Arrow (v2.3.3)[26] with the PacBio subreads. The FALCON assembly yielded 3,656 contigs with an N50 and mean length of 4.66Mb and 677Kb respectively (Fig.S10). FALCON contigs were scaffolded with chromium linked reads by ARCS (v1.05)[26] and LINKS (v1.8.6)[27]. The link ratio (-a, default 0.3) was set to 0.9 as recommended for a scaffolding application. 1,170 Falcon contigs were joined in this step, which increased the scaffold N50 of the assembly to 18.5Mb.

To evaluate the correctness of the scaffolding, we aligned the assembled sequences on a high-density canine linkage map[28]. 21,278 reported markers were unambiguously mapped to the assembly by BLAT (v36)[29]. The synteny of genetic and physical location of markers were further compared by Chromonomer (v1.08)[30]. In this step, 207 scaffolds were anchored on the linkage map, and four scaffolds were reported as having conflicting markers. After carefully reviewing the sequence, we confirmed that all discrepancies came from incorrect joining of sequences from different chromosomes and therefore split these four scaffolds.

We then performed gap filling using the PacBio subreads (PBjelly from PBSuite v15.8.24[31]), and 648 gaps were closed during this process. The assembly was further scaffolded by HiC (HiRise, Dovetail Genomics). Before scaffolding, an initial QC scan was performed using the HiC linked reads on the input sequence and no putative wrong joins were reported. With the long-distance interaction information from HiC, the assembly was successfully scaffolded to chromosome-level (scaffold N50: 64.3Mb). To evaluate the HiC results, the JUICER[32] pipeline was used to map the HiC reads back to the HiRise assembly to generate and visualize a HiC map with intra- and inter-chromosomal interactions. Based on this map, we identified and manually adjusted contigs placed in the wrong order or orientation on 5 chromosomes (chr6, 14, 17, 26 and X), and joined separated contigs from the same chromosome (chr8 and 18). A second round of gap filling was performed by PBjelly and 110 gaps were closed. To improve the accuracy, we polished the assembly by Arrow with PacBio subreads and also Pilon (v1.22)[33] with the 10x Genomics reads (BWA mem, v0.7.15[34]), respectively. We applied a FreeBayes-based method to correct the remaining Indel errors in the assembly[35]. We aligned the short reads to the polished assembly, and called the SNPs and Indels by FreeBayes (v1.1.0)[36] with setting “-C 2 -0 -O -q 20 -z 0.10 -E 0 -X -u -p 2 -F 0.75”. We replaced the reference base with the variant allele at 149,264 positions where 10x Illumina sequencing depth was at least 30X and and the variant allele ratio was >90% using FastaAlternateReferenceMaker from GATK (v4.1.1.0)[37]. A final round of Pilon short-read polishing was performed after the allele replacement. We conducted taxonomic classification (Kraken2, v2.0.8[38]) for the assembly. A total of 68 unplaced contigs were removed because they were suspected to be from bacterial contamination.

The assembly is available at NCBI (DDBJ/ENA/GenBank: JAAHUQ000000000), and UCSC/Trackhub (https://genome-test.gi.ucsc.edu/cgi-bin/hgTracks?db=canFam4).

### Mapping the end-sequences of BAC clones from CH82

End sequences from bacterial artificial chromosome (BAC) clones in the CH82 library were extracted from the TraceDB of NCBI (ftp://ftp.ncbi.nih.gov/repository/clone/reports/Canis_familiaris/CH82.endinfo_9615.out). The end sequences were mapped as paired-reads by BWA mem, with default setting, on GSD1.0 and CanFam3.1. Only end pairs that could map on both assemblies were selected for the analysis. The end pairs were defined as concordant pairs when they were aligned in forward and reverse direction with a distance <500Kb.

### Assembly QC and properties

We assessed the GC-content of the GSD1.0 by scanning the assembly using a 50bp window, and percentages of guanine and cytosine in each window was calculated with NUC from BEDTools (v2.29.2)[39]. CpG islands were detected by the “cpg_lh” script from UCSC utilities (http://hgdownload.soe.ucsc.edu/admin/exe/linux.x86_64.v369/), with a modified method described by Gardiner-Garden[40]. We assessed the mappability on GSD1.0 with different k-mer of 50/150/250bp using GEM-Tools (v1.71)[41].

Repetitive elements including transposable elements (SINE/LINE/LTE/DNA elements) and tandem repeats were annotated by Repeat Masker (v4.0.8, http://www.repeatmasker.org) in a sensitive mode with a combined library (dc20171107-rb20181026). Specifically, to identify telomere sequences, we searched for the “TTAGGG” repeat on both the plus and minus strands of GSD1.0 by fuzznuc from EMBOSS (v6.6.0)[42]. A putative telomere sequence was defined as to have at least 12 consecutive repeats with less than 11 variant bases between the repeats. Candidate telomeric repeats within 100bp were merged. The detection of centromere was performed by scanning the assembly with 5kb windows to calculate the percentage of three satellite repeats (CarSat1/ Carsat2/ SAT1_CF), known to be associated with centromeres[43]. Any window with content of these three repeats >80% was considered as the putative centromere sequence.

### Gap closure in GSD1.0 compared to CanFam3.1

Any continuous ambiguous “N” bases in CanFam3.1 were considered as a gap. For each gap, we extracted the 1kb flanking sequences, and mapped these as pairs to GSD1.0 with BWA mem. The gap of CanFam3.1 was considered as closed when 1) flanking sequence pairs could be mapped properly in same scaffold with mapping quality >20; 2) the distance between the pairs was less than 100 kb; and 3) no gap in GSD1.0 was present in the sequence between pairs. With this approach, we could identify the sequence for 18,649 of 19,553 (95.4%) gaps from assembled chromosomes and 1,563 of 4,323 (36.2%) gaps from unplaced scaffolds of CanFam3.1 in GSD1.0. The flanking sequences of 3,072 gaps overlapped each other in GSD1.0, suggesting an artificial gap in CanFam3.1. However, these regions can be considered closed in GSD1.0. For the other closed gaps, we extracted the filled sequences from GSD1.0 and calculated GC and repeat content. BEDTools was used to intersect exons, miRNA and ATAC-seq peaks with those filled gaps. Specifically, we looked for novel genes from the filled gaps. A novel gene was defined if it 1) had at least 80% of gene body identified from the filled gaps; 2) was not a pseudogene; 3) had not been annotated in the unplaced scaffolds of CanFam3.1; 4) did not have the duplicated/homologous fragment in another region of the genome. With these thresholds, we found 8 novel genes from the filled gaps, and all locate in the regions with good synteny of human hg38 assembly.

### RNA preparation and long-read cDNA sequencing

Multiple Beagle total RNA and tissue samples for full-length cDNA and miRNA sequencing experiments were sourced from Zyagen (USA; Additional file 2: Tab. S10). Total RNA from Beagle hypothalamus (RIN>8) was used for PacBio Iso-Seq following the Iso-Seq express protocol. Two libraries were made and each was sequenced on a single SMRT-cell on the Sequel system, yielding approximately 500,000 reads each with mean read lengths of 2,452 bp and 3,451 bp.

For nanopore and miRNA-sequencing, samples from either a male or a female Beagle were used to prepare total RNA using a standard TRIzol protocol (Invitrogen, US). In total, libraries from 25 tissues were sequenced, with a focus on different brain regions (Additional file 2: Tab. S11). The PCR strand-switch protocol and the SQK-LSK109 kit were used for MinION sequencing (Nanopore, UK). Zyagen samples were amplified with PBC096 barcoding for 8-10 cycles with both LongAmp (female, 62°C annealing; NEB, US) and PrimeSTAR GXL (male and female, 64°C annealing; Takara bio, US), with a 10 minutes extension time. A separate Beagle retina sample was sequenced using both the nanopore direct cDNA sequencing kit SQK-DCS109 and on an NovaSeq 6000 S4 lane (Illunima, US). Reads were base called with the high accuracy model in guppy (v3.6 for direct cDNA and v3.3 for amplified samples). After barcode demultiplexing with Qcat (https://github.com/t-neumann/qcat) we used pychopper (Nanopore, UK) to identify and orient fully sequenced reads. To improve mapping accuracy only reads with a quality value above 15 was used. Nanopore cDNA sequences were basecalled with the high accuracy R9.4.1 model in Guppy (v3.3) and processed with pychopper to identify and orient full-length reads. Only reads with average base quality score above 15 were used. For PacBio, full-length CCS reads with at least three passes were selected. The long-read cDNA runs were mapped with Minimap2[44] v2.17 with the options - x splice -G 500000 and --junc-bed with splice junctions identified from the Illumina alignments. These settings improved mapping to genes with long introns and short exons.

MicroRNA libraries were made with the NEXTFLEX small RNA library kit v3 (PerkinElmer, US) and 25 million reads were generated with a NextSeq500 instrument (75bp high-output kit v2.5 in paired-end mode).

### Gene annotation

We searched for dog RNA-seq sequences in NCBI using SRA-Explorer (https://sra-explorer.info/) aiming to select read sets from a diverse set of tissues and breeds and included stranded samples with paired reads of at least 100bp (Additional file 2: Tab. S11)[42–45]. Reads from the same study and tissue were combined and adaptors were trimmed with BBmap. RSeQC was used on a small subset of reads for each sample to infer library type and the corresponding setting was used for alignment with HISAT2[45]. Illumina reads were processed using the superreads module in Stringtie2[46] to improve mapping. Transcripts were assembled with settings -f 0.05 as the threshold for isoforms expression and the gtf-files were merged with stringtie --merge. Assembled transcripts were processed with TAMA tools[47] for ORF detection and BLAST parsing to identify coding regions. We used curated proteins from Uniprot_Swissprot together with proteins from the latest ENSEMBL dog annotation (v100) and selected the longest blast hit from the top 5 hits with an E-value below 10^-10 as the id of the protein. Long reads were assembled both with Stringtie2 in combination with the short reads, and separately with TAMA tools. For the final annotation, bam files for the same tissue type were merged before stringtie assembly, and additional long read transcripts from TAMA were added if they either had a blast hit >90% or were multi-exonic, located outside of the stringtie annotation and had a blast hit covering >50% of the target. Additional filtering was made to remove transcripts that 1) were long single exon transcripts (>10kb and <10% intronic sequence) or 2) originated from genomic polyA/T regions. Gffread was used to re-group transcripts into genes, and only one transcript per unique CDS region was kept. Finally, transcripts with a bad BLAST classification (<50% hit) were removed if they belong to a group with high scoring transcripts (Additional file 1: Fig. S11).

Micro-RNA-seq samples were downloaded from SRA and combined with our brain miRNA-seq reads for a total of 1.3 billion reads. After adaptor trimming and collapsing of unique sequences between 20 and 30 bases, MiRDeep2[48] was used to identify micro-RNAs with mirBase entries for dog and human as comparison. ATAC-seq reads from BARKbase[49] were aligned with BWA mem and peaks called with Genrich (https://github.com/jsh58/Genrich). BedGraph files were produced with BEDTools.

### Assembly benchmark with Busco and Iso-Seq data

We ran BUSCO (v3.0.2b) with the mammalia_odb9 dataset which includes 4,104 single copy genes that are evolutionarily conserved between mammals to assess the completeness of the assemblies. To ensure reads with high sequence base accuracy, only PacBio circular consensus sequencing (CCS) reads supported by >10 subreads were used for the benchmarking (277,280 reads from s003 and 256,422 reads from s004). The high-quality CCS reads were mapped to GSD1.0 and CanFam3.1 by the minimap2 (v2.17, “-x splice:hq -uf -t 8 --cs”). We calculated the percentage of mapped bases for each read according to the “difference string” in cs tag. With the same method, we also benchmarked five newly released canine assemblies (Additional file 2: Tab. S3), including the Luka (Basenji), Nala (German Shepherd), Zoey (Great Dane), Scarlet (Golden Retriever), and the Sandy (Dingo) assembly.

### 10x and standard Illumina short read (ISR) mapping

In order to investigate the genetic variations across the breeds, we also prepared HMW DNA from 27 dogs (19 breeds) and sequenced them with Illumina HiSeq X (2×150 bp). The sequencing depth ranged between 34-110X, with an average depth of 46X and the highest depth for the Mischka reference sample (Additional file 2: Tab. S6).

We mapped the Chromium reads to GSD1.0 with the Long Ranger (v2.2.2, 10X Genomics, USA) WGS pipeline. Due to a limit of scaffold numbers in Long Ranger, we concatenated all unplaced scaffolds of GSD1.0 into a single scaffold with 500 “N” bases between scaffolds. The aligner in Long Ranger, Lariat, maps Linkedeads with the same barcode simultaneously in a given genomic range according to the input molecule length, which increases the mapping quality of reads from duplicated regions.

SNPs and short Indels were detected in 10x and ISR dataset using different modules from GATK4. Variants were called from alignment by HaplotypeCaller, and further merged by the CombineGVCFs and GentoypesGVCFs. The SNPs and Indels were filtered by SelectVariants with "QD < 2.0 || FS > 60.0 || MQ < 40.0 || MQRankSum < −12.5 || ReadPosRankSum < −8.0" and "QD < 2.0 || FS > 200.0 || ReadPosRankSum < −20.0", respectively.

### Dark regions detection

To study the dark regions for different sequencing technologies, we downloaded the Illumina short read (ISR) data of 25 dogs from public resources (Additional file 2: Tab. S12), with breeds selected to match the dogs sequenced with 10x. Read pairs were mapped to the GSD1.0 using BWA mem with default settings. For each sample, the coverage was calculated by bamCoverage from Deeptools (v3.3.2)[50] with a 25bp window, whereas unmapped reads and secondary alignment were excluded from the analysis. Meanwhile, we also assessed the proportion of reads with mapping quality >10 in each window. To identify dark regions by coverage (COV), we searched the genomic windows with coverage ≤5X. However, this threshold was adjusted for sequencing depth; a lower cutoff was applied in low coverage samples to select the ~60Mb genomic regions (Additional file 2: Tab. S13). The discovered individual COV dark regions were merged, and the COV fraction for each window was assessed in two ISR and 10x datasets: windows with F_COV_ >0.9 (90% individuals, in at least 23 ISR dogs or 25 10x dogs) remained as the candidate COV dark regions.

Regions were defined camouflaged (CAM) if the coverage was ≥10X and the proportion of high mapping quality reads was less than 10%. We searched for and merged the genomic windows that reached the threshold from each dog. Notably, the CAM regions detected in one individual could have been assigned as COV in the others. Thus, we excluded those COV dark dogs before we calculated the fraction of CAM for each window; any window with F_CAM_ >0.9 was selected as a candidate.

### Structural variation detection

In order to build a high-resolution map of structural variation, we scanned the genomes of 27 10x dogs using four SV callers. First, 1) Long Ranger, which was used to call the SVs in two size ranges. (a) Smaller SVs ranged from 50bp to 30Kb, with Long Ranger examining the haplotype-specific coverage drops and discordant reads pairs. (b) Large-scale SVs > 30Kb, where Long Ranger detected the paired coverage of genomic loci that shared many more barcodes than expected by chance. Candidate SVs were further refined and categorized by comparing the layout of reads and barcodes around the breakpoints. Three additional callers were adapted to discover other types of median size SVs (50bp-30kb). 2) GridSS[51] and 3) manta[52] are assembly-based callers which have been reported to have a good performance in different studies[53,54]. Both detected SVs using evidence from split and paired reads, and also assembled the sequences of breakpoints to accurately estimate these positions. The type of SVs called by GridSS was determined by the orientation of reads from the breakpoints using a R script (https://github.com/PapenfussLab/StructuralVariantAnnotation). From the three callers above, only high-quality SV calls marked as “PASS” in vcfs were kept for analysis. Lastly, 4) CNVnator[55], predicted the copy number variations by a read-depth (RD) approach. A 150bp bin size was suggested for screening, and called SVs were QC filtered by requiring a p-value <0.05 for a RD t-test statistic (e-val1) and the probability of RD frequency <0.05 in a gaussian distribution of (e-val2). The result was converted into VCF form using the “cnvnator2VCF.pl” script from the CNVnator package.

For each dog, the filtered median SVs from all four callers were merged by the SURVIVOR[56], and combined with the large size SVs called from Long Ranger. Specifically, we removed the SVs on chromosome X that were only supported by CNVnator, since the algorithm lacks the right model for SV detection for chromosome X in females. Finally, SVs were further merged across individuals into a nonredundant SVs set.

### SV validation and genotyping

SVs for validation were selected based on their position overlapping protein-coding genes, and on being polymorphic in the 10x data set (>3/27 individuals contained the structural variant).

Genomic DNA was extracted from whole blood using either MagAttract HMW DNA Kit (QIAGEN, Germany) or NucleoSpin Blood Kit (Macherey-Nagel, Germany). SV breakpoints were confirmed with Sanger Sequencing where possible. PCR was performed with either PrimeSTAR GXL DNA Polymerase (Takara, US) or AmpliTaq Gold DNA Polymerase (ThermoFisher, US) according to manufacturer's recommendations. PCR fragments were cloned using either Zero Blunt or TOPO TA Cloning Kit (Invitrogen, US) depending on PCR overhang. Plasmid DNA was extracted using QIAprep Spin Miniprep Kit (QIAGEN, Germany), PCR products and plasmids were sequenced using the Mix2Seq service (Eurofins Genomics, Germany) and analysed using CodonCode Aligner v6.0.2 (CodonCode, US).

For deletion PCR genotyping, reactions were performed as single or multiplex reactions depending on variant size, and visualised using 2% TAE agarose gels. For CNV genotyping, ddPCR absolute quantification was performed using 15 ng of *DraI* or *AluI* (NEB, US) pre-restriction digested DNA in a 20ul reaction mix containing 1x ddPCR Supermix for Probes (BioRad, US) with 900nM target and reference primers (Integrated DNA Technologies) and 250nM of target and reference probes (Thermo Fisher Scientific, US). Reactions were quantified using a QX100 instrument and analysed with QuantaSoft v1.3.1.0 (BioRad, US).

All primers and probes were designed using Primer3 v0.4.0 (http://bioinfo.ut.ee/primer3-0.4.0/) and collated in Tab. S8 (Additional file 2).

### Gene Expression Analysis

Total RNA was extracted from liver, spleen and heart tissues using the AllPrep DNA/RNA/miRNA Universal Kit (QIAGEN, Germany) according to manufacturer's specification and including on-column *DNaseI* treatment (Additional file 2: Tab. S14). 500-1000ng of total RNA was reverse transcribed using the Advantage RT-for-PCR Kit (Takara) and qPCR performed in quadruplet using SYBR Green PCR Master Mix (Thermo Fisher Scientific, US) and 900nM primers in a QuantStudio 6 Real-Time system (Thermo Fisher Scientific, US) with standard cycling and dissociation curve analysis. Two housekeeper primer sets (RPS19 and RPS5) were assessed for stability (*Normfinder* R package[57]) and used in combination to calculate relative gene expression[58]. These calculations included primer specific efficiencies and used the average Ct from all control samples for initial delta Ct normalisation. wilcox.test in R was used to assess the significance of between genotypic class gene expression changes.

## Supplementary information

Additional file 1: supplementary figures

Figure S1. Genetic position of Mischka using the 170K SNP chip.

Figure S2. Sequence characteristics of 2,159 unplaced scaffolds in GSD1.0.

Figure S3. Tree map of contig sizes of CanFam3.1 and GSD1.0.

Figure S4. Content of repetitive element on each chromosome.

Figure S5. Iso-Seq data mapped to GSD1.0 and CanFam3.1.

Figure S6. Comparison between length of the best BLAST hit for genes in GSD1.0 and CanFam3.1 annotation.

Figure S7. Mirlet7i identified from GSD1.0.

Figure S8. Closure of gaps in cancer genes from COSMIC.

Figure S9. Sequence comparison of immunity loci between GSD1.0 and CanFam3.1.

Figure S10 Illumina short reads (ISR) dark regions rescued by 10x sequencing.

Figure S11. RNA sequencing of different tissues.

Additional file 2: supplementary tables

Table S1 Assembly statistics for GSD1.0 and CanFam3.1

Table S2 Summary of repetitive elements in GSD1.0

Table S3 Summary of available canine assemblies from public resources

Table S4 ISO-seq reads mapping in different canine assemblies

Table S5 Filled gaps within the cancer genes from COSMIC

Table S6 10x sequencing of Mischka and 27 dogs

Table S7 Primers used for validation and genotyping

Table S8 Gene expression summary for structural variant loci.

Table S9 Sequencing data generated from three HiC libraries for Mischka dog

Table S10 Tissues used for nanopore sequencing

Table S11 Datasets used for annotation

Table S12 Public resources of illumina short-read data of 25 dogs

Table S13 Summary of dark regions detected in each dog

Table S14 Tissues samples and genotyping results for expression analysis

## Abbreviations

## Declarations

### Ethics approval and consent to participate

Approval was obtained from dog owners before collecting the biological samples at veterinary clinics. Ethical approvals for sampling were granted by Uppsala Animal Ethical Committee and Swedish Board of Agriculture (C417/12, C418/12, C103/10, C12/15 and C15/16). Importation of canine tissues was approved by Jordbruksverket (6.7.18-14513/17).

### Consent for publication

Not applicable

### Availability of data and materials

The PacBio-long reads, HiC, and Illumina 10x data of Mischka are available in SRA under BioProject PRJNA587469. The Illumina 10x data of 27 dogs are available in SRA under BioProject PRJNA588624. Scripts used in the study is available at the GitHub repository (https://github.com/Chao912/Mischka/). The canFam_GSD1.0 assembly is deposited in DDBJ/ENA/GenBank under JAAHUQ000000000.

## Authors’ contributions

KLT, JRSM and MLA conceived the study and designed the experiments. GP and MLA collected the samples with the help of JH1, ÅO, SS, HR, IL, SM, JH2 and ÅH. MLA, ÅK and ÅO performed the DNA/RNA extractions. ÅK, ES and JRSM performed the validation of structural variation, genotyping and expression analyses. CW, OW, MLA and KLT contributed to the data analysis of the genome assembly. OW performed the gene annotation with the help of TB and SM. JRSM and KLT oversaw and interpreted the results together with CW, OW and MLA and ES. CW, OW, JRSM and KLT wrote the manuscript with input from all authors.

## Acknowledgements

We thank Mischka’s owners who kindly allowed us to collect blood and tissues for scientific purposes. We would like to acknowledge Mats Pettersson, Olga Vinnere Pettersson and Ignas Bunikis for helpful suggestions. We thank SNIC through Uppsala Multidisciplinary Center for Advanced Computational Science (UPPMAX) for providing computation resources under Project SNIC 2017/7-384, 2017/7-385 and 2020/5-190. Sequencing was performed by the SNP&SEQ Technology Platform in Uppsala. The facility is part of the National Genomics Infrastructure (NGI) Sweden and Science for Life Laboratory. The SNP&SEQ Platform is also supported by the Swedish Research Council and the Knut and Alice Wallenberg Foundation.

