## additional file 1 for "A new long-read dog assembly uncovers thousands of exons and functional elements missing in the previous reference"

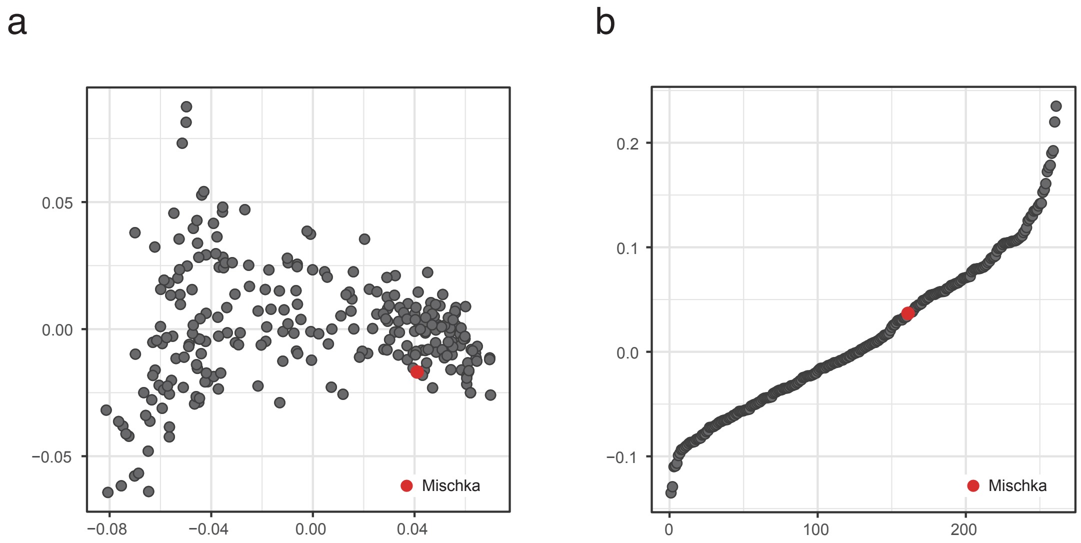


**Figure S1. Genetic position of Mischka using the 170K SNP chip.** We genotyped Mischka with the 170K Illumina SNP chip, and merged the data with 260 Swedish German Shepherds from a previous study[24]. (a) Multidimensional scaling (MDS) plot shows that Mischka falls within the cluster of other German Shepherds. (b) The inbreeding coefficient for Mischka is 0.037, indicating that Mischka has an acceptable inbreeding level.

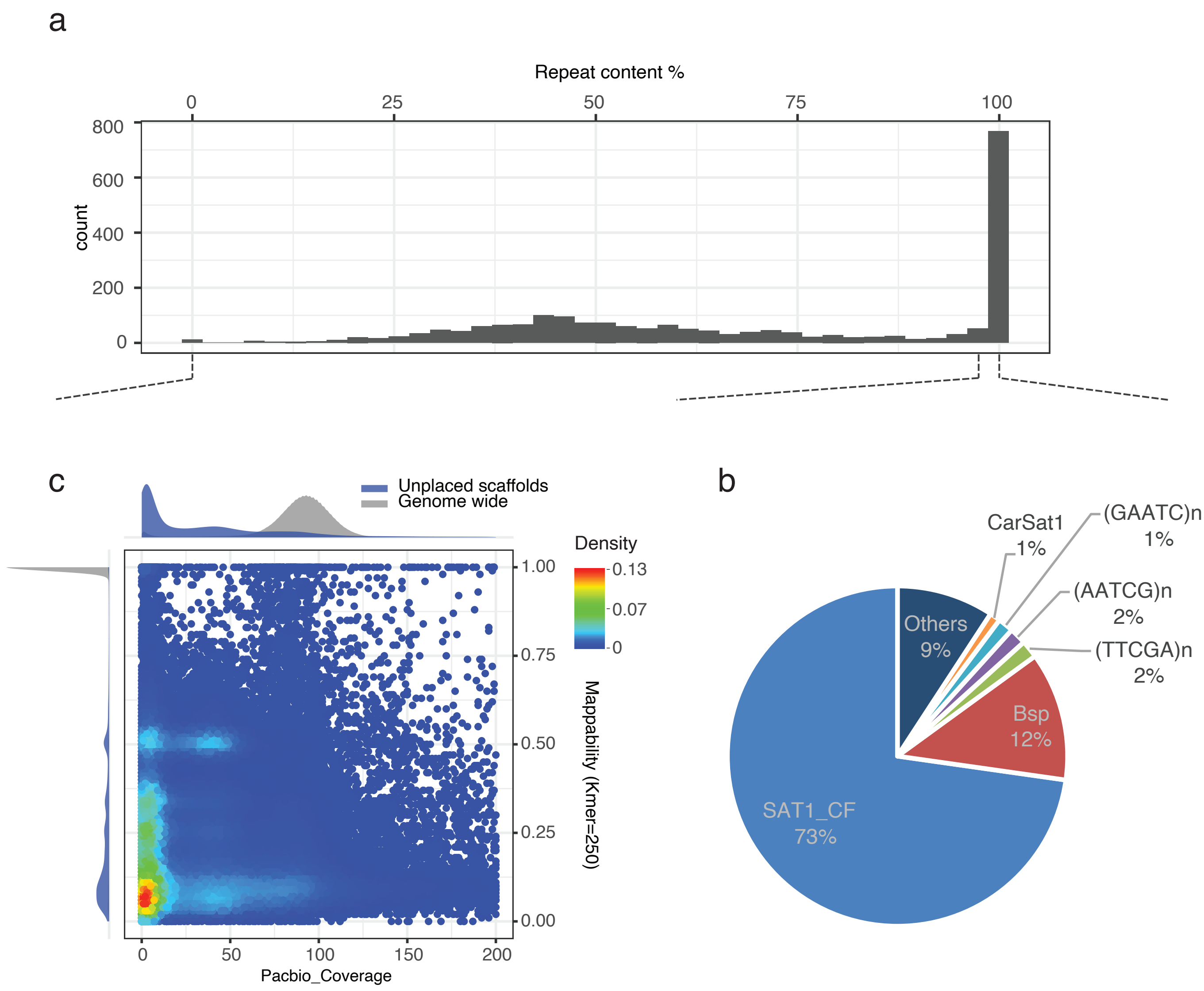


**Figure S2. Sequence characteristics of 2,159 unplaced scaffolds in GSD1.0.** For each unplaced scaffold, we calculated the repeat content and the average of mappability (k-mer = 250bp). (a) Distribution of the repeat content for unplaced scaffolds. 871 scaffolds are found with content >90%. (b) The most common repeat is SAT1_CF, which is known as a centromeric repeat. (c) Of the remaining scaffolds, most have lower coverage and mappability compared to the chromosome level scaffolds, suggesting they could be from the genomic duplication regions.

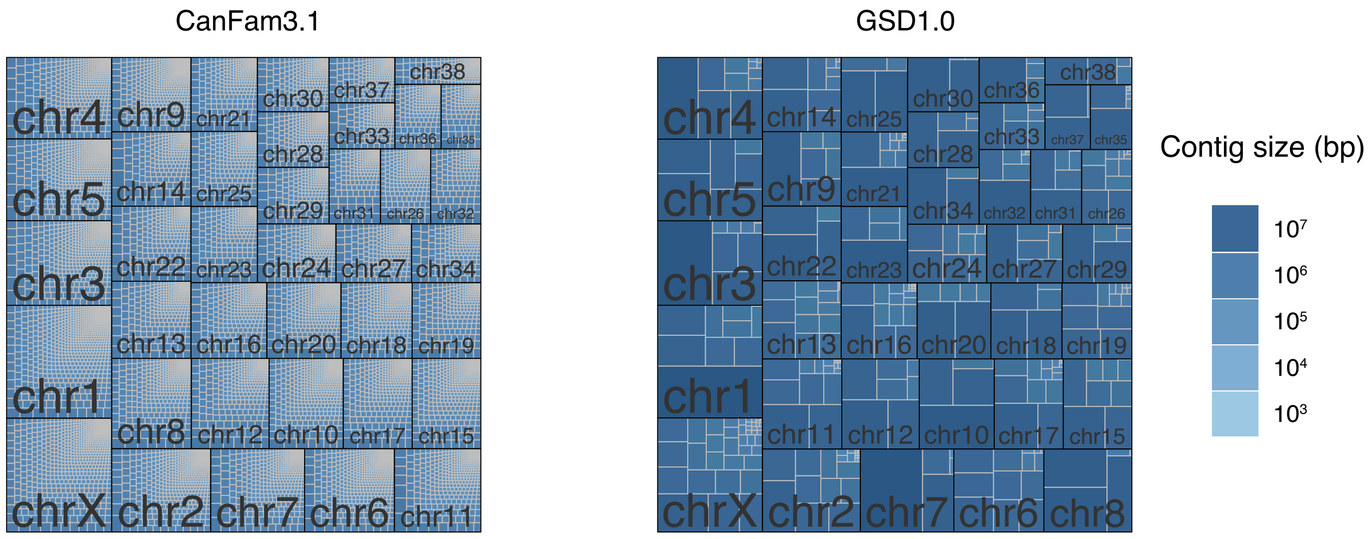


**Figure S3. Tree map of contig sizes of CanFam3.1 and GSD1.0.**

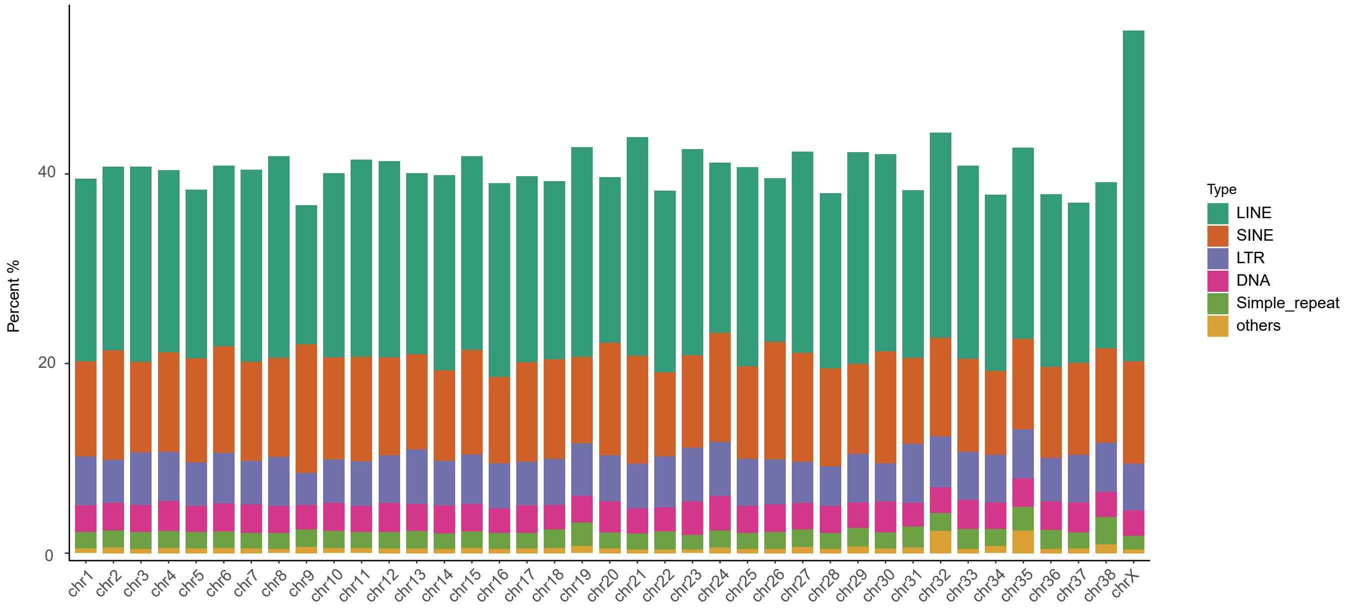


**Figure S4. Content of repetitive element on each chromosome.** The repeat-content is stable between autosomes. Chromosome X shows a relatively higher repeat content by having more LINEs.

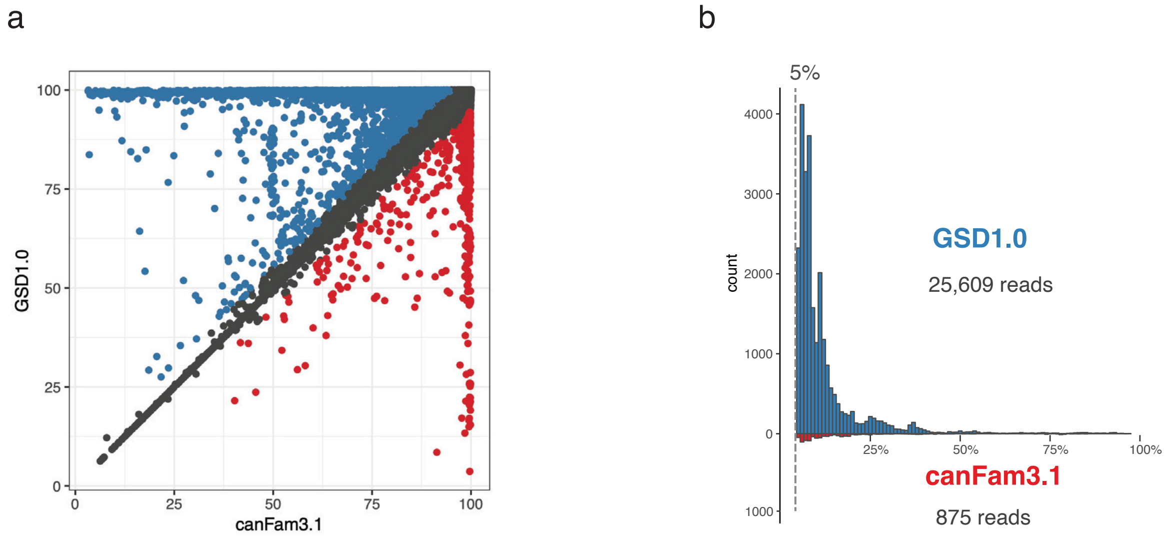


**Figure S5. Iso-Seq data mapped to GSD1.0 and CanFam3.1.** (a) The percentage of mapped bases for each read in the two assemblies. Each dot indicates an Iso-Seq read. Blue dots showed the reads with >5% more bases mapped in the GSD1.0 than CanFam3.1. Red dots indicated the reads have 5% more bases mapped in CanFam3.1. b) Distribution of mapping improvement for two kinds of reads. Altogether 25,609 reads have at least 5% more bases mapped in the GSD1.0, whereas only 875 reads could be better mapped in CanFam3.1.

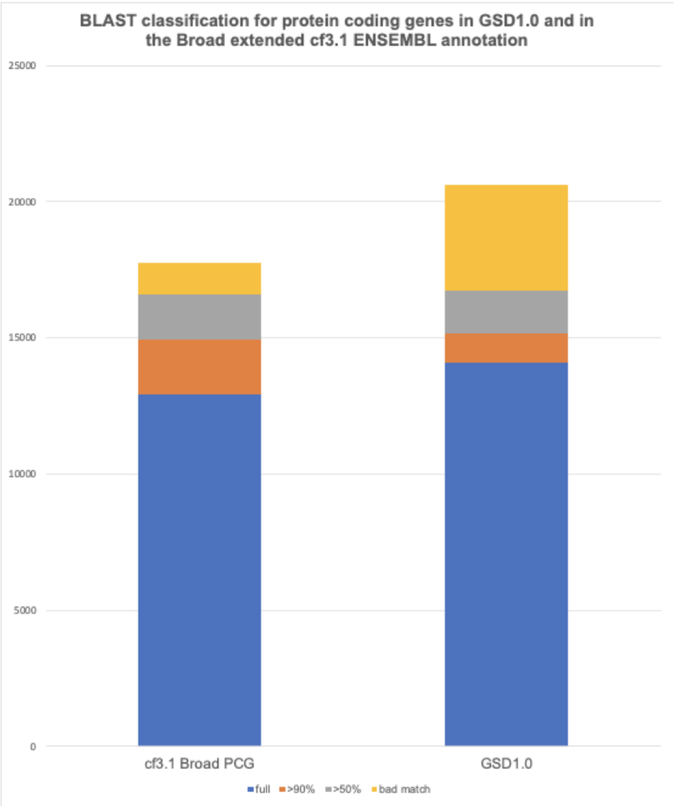


**Figure S6. Comparison between length of the best BLAST hit for genes in GSD1.0 and CanFam3.1 annotation.**

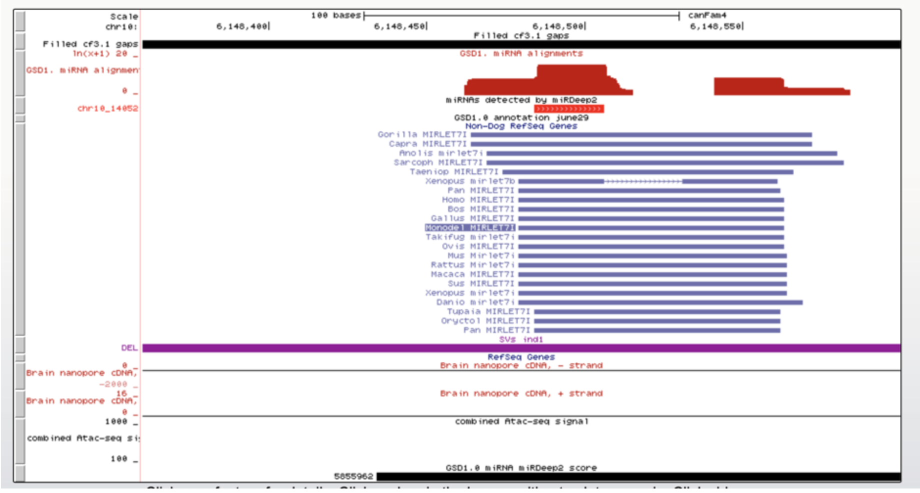


**Figure S7. Mirlet7i identified from GSD1.0.** Mirlet7i has the 16th highest miRDeep2 score, with close to 11 million aligned reads from the combined miRNA-seq dataset within a filled gap.

a

b

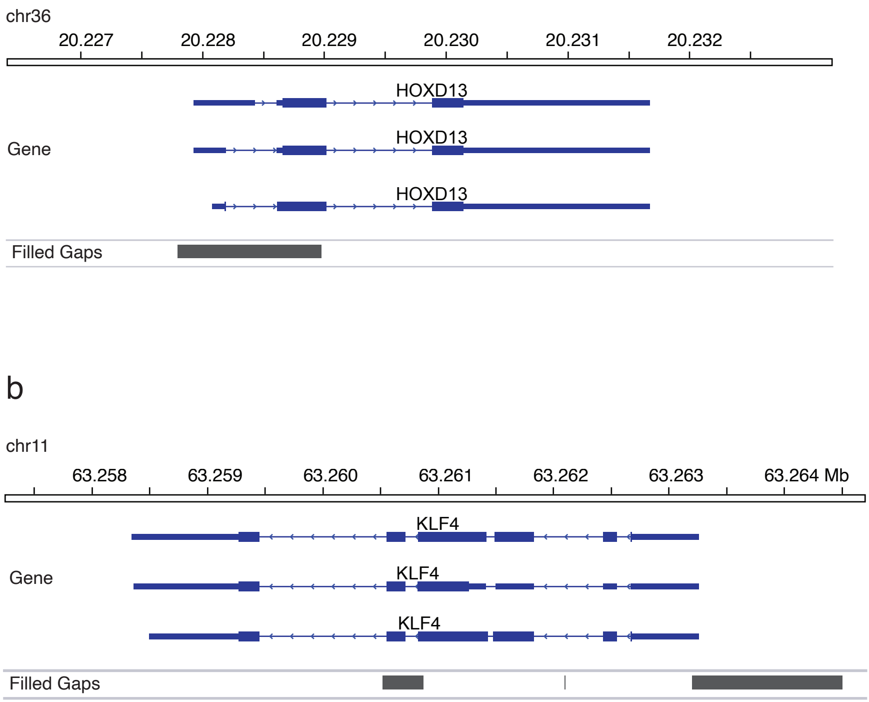


**Figure S8. Closure of gaps in cancer genes from COSMIC.** (a) *HOXD13* gene and (b) *KLF4* gene

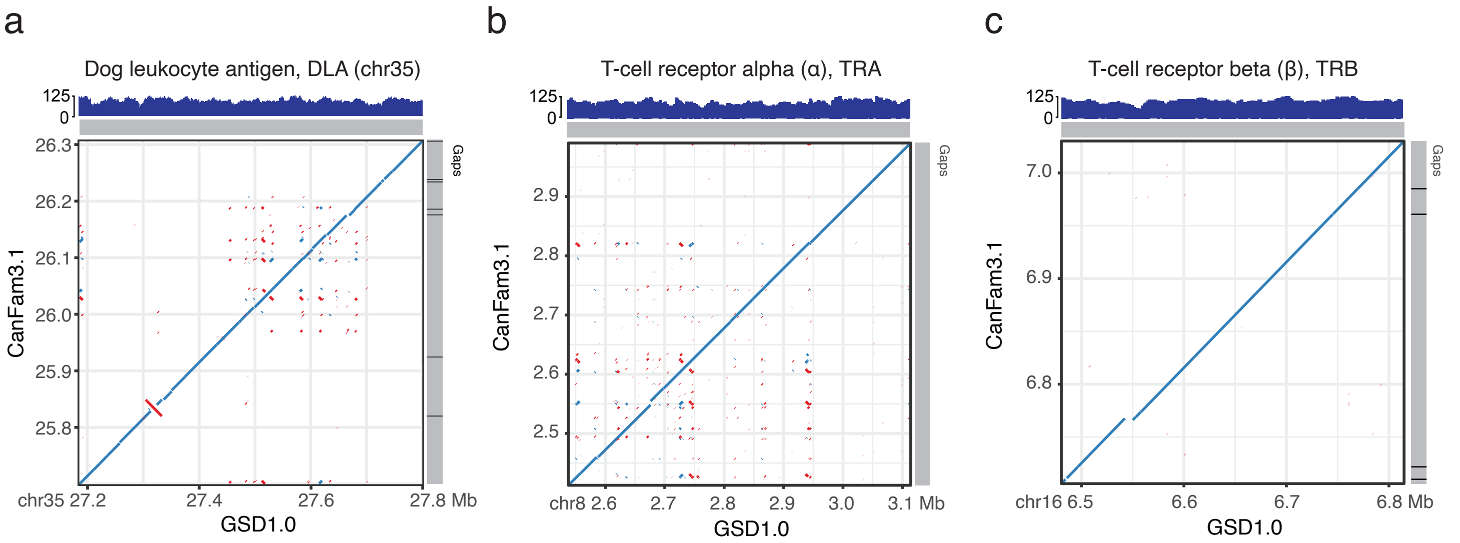


**Figure S9. Sequence comparison of immunity loci between GSD1.0 and CanFam3.1.** (a) Dog leukocyte antigen locus (DLA) on chromosome 35. (b) The T-cell receptor alpha (TRA) locus on chromosome 8, and (c) T-cell receptor beta (TRB) locus on chromosome 16.

a

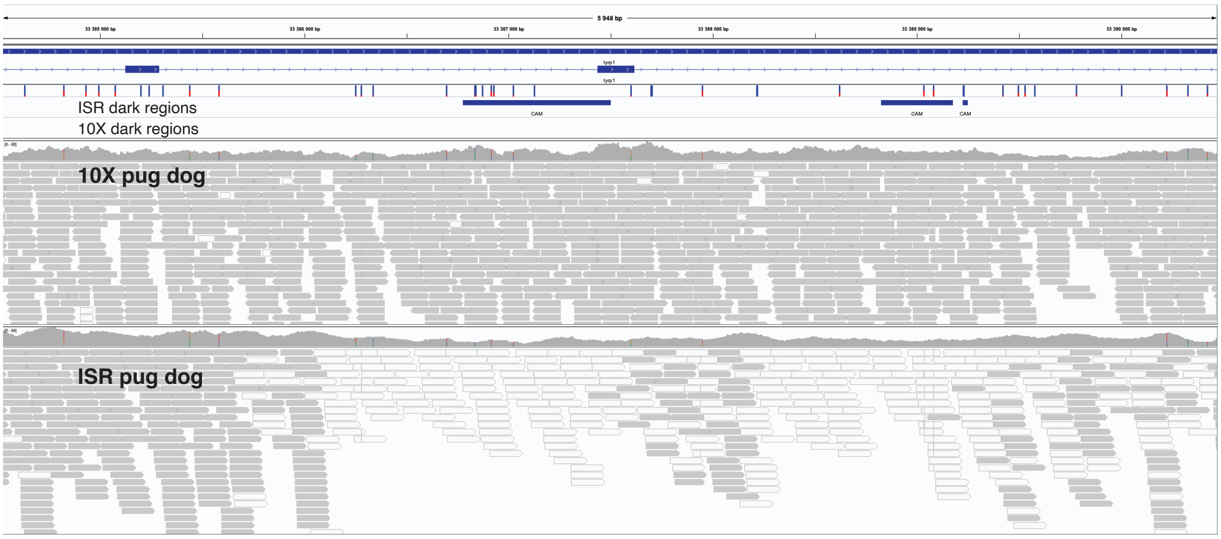


b

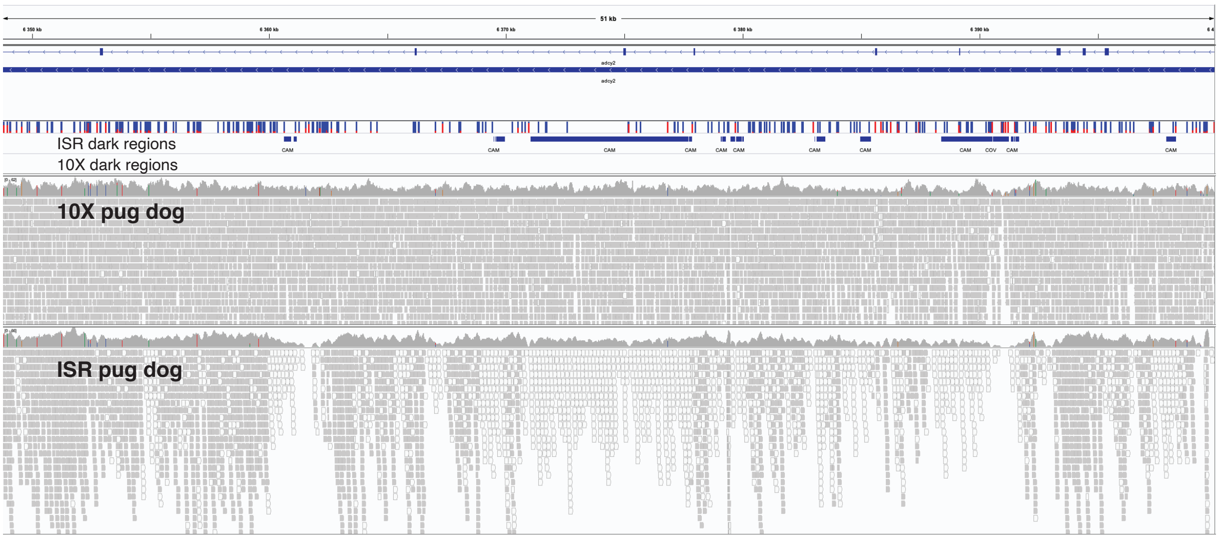


**Figure S10 Illumina short reads (ISR) dark regions rescued by 10x sequencing.** IGV snapshot shows the alignments of a pug dog sequenced by Illumina short reads and 10x linked reads. a) ISR dark regions in the *TYRP1* gene. b) ISR dark regions in the *ADCY2* gene. These ISR dark regions have been recovered by 10x linked reads sequencing, where the reads showed high mapping quality.

a

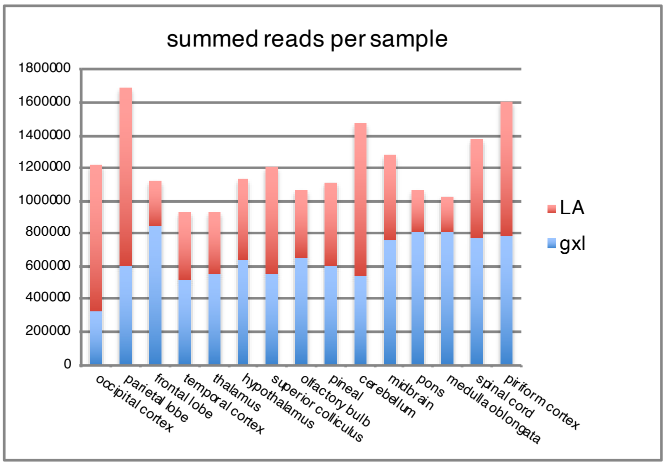


c

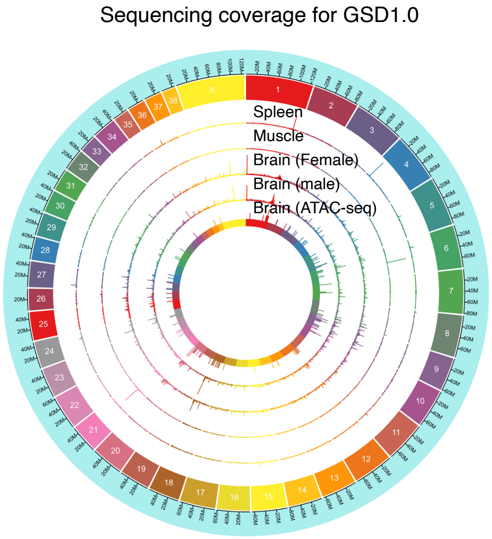


b

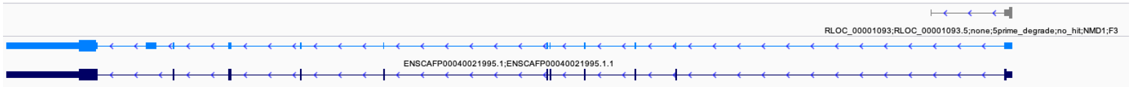


**Figure S11. RNA sequencing of different tissues.** (a) Total read numbers per tissue for a multplexed run (2 flowcells) with female brain samples. N50 of reads was 2.4 kb, and amplification was done with two enzymes (LongAmp and GXL) to avoid bias in coverage. (b) Annotation were to remove degraded transcripts (gray). Colors indicate the quality of BLAST hit as defined by TAMA: gray=no hit, dark blue= full hit, medium blue=90% match, light blue=50% match, light skye blue=bad hit. (c) Read distribution for Nanopore samples for various tissues. To avoid issues with high coverage from tissue specific genes in transcript assembly and merging reads were assembled per sample type and resulting transcripts merged.
