## additional file 2 for "A new long-read dog assembly uncovers thousands of exons and functional elements missing in the previous reference"

Table S1 Assembly summary of the GSD1.0 and CanFam3.1

| | GSD1.0 | CanFam3.1 |
|---|---|---|
| number of contigs | 2,783 | 27,104 |
| N50 (L50) contig | 14,840,767 bp (57) | 267,478 bp (2,436) |
| Number of scaffolds | 2,198 | 3,268 |
| N50 (L50) scaffolds | 64,299,765 bp (15) | 63,241,923 bp (15) |
| Number of Gaps | 585 | 23,876 |
| Gap density (gaps/Mb) | 0.24 | 9.9 |
| Total bases | 2,482,000,080 bp | 2,410,976,875 bp |
| total ungaped bases | 2,481,941,580 bp | 2,392,715,236 bp |

Table S2 Summary of repetitive elements in GSD1.0

| Type<br>subtype | number of elements | length occupied (bp) | percentage of sequence (%) |
|---|---|---|---|
| SINEs | 1538330 | 253126533 | 10.2 |
| Alu/B1 | 0 | 0 | 0 |
| MIRs | 436540 | 64619318 | 2.6 |
| LINEs | 878624 | 504530620 | 20.33 |
| LINE1 | 534540 | 413420208 | 16.66 |
| LINE2 | 292578 | 79467264 | 3.2 |
| L3/CR1 | 37964 | 8374531 | 0.34 |
| RTE | 12346 | 3077052 | 0.12 |
| LTR element | 308696 | 120262262 | 4.85 |
| ERVL | 90609 | 40692875 | 1.64 |
| ERVL-MaLRs | 146071 | 50536666 | 2.04 |
| ERV_classI | 49789 | 23477942 | 0.95 |
| ERV_classII | 0 | 0 | 0 |
| DNA element | 325449 | 68706785 | 2.77 |
| hAT-Charlie | 187859 | 36590816 | 1.47 |
| TcMar-Tigger | 49675 | 14764477 | 0.59 |
| Unclassified | 5213 | 965509 | 0.04 |
| Total interspersed repeats | | 947591709 | 38.18 |
| Small RNA | 1141119 | 191329823 | 7.71 |
| Satellites | 3922 | 52090599 | 2.1 |
| Simple repeats | 955316 | 50351697 | 2.03 |
| Low complexity | 129088 | 7138943 | 0.29 |

Table S3 Summary of available dog assemblies from public resources

| Assembly | Origin | Dog name | Sequencing Tech. | total length (bp) | Number of scaffolds | N50 (bp) | NCBI Accession/Links |
|---|---|---|---|---|---|---|---|
| CanFam3.1 | Boxer | Tasha | BAC+Sanger sequencing | 2,410,976,875 | 3,310 | 45,876,610 | GCF_000002285.3 |
| CanFam GSD1.0 | German Shepherd | Mischka | Pacbio+10X+HiC | 2,482,000,080 | 2,198 | 64,299,765 | GCA_011100685.1 |
| ASM864105v3 | German Shepherd | Nala | Pacbio+BioNano+HiC | 2,407,242,830 | 410 | 64,346,267 | GCA_008641055.3 |
| UMICH_Zoey_3.1 | Great Dane | Zoey | PacBio | 2,343,218,756 | 794 | 64,204,256 | GCA_005444595.1 |
| ASM325472v1+HiC | Dingo | Sandy | Pacbio+BioNano+10X+HiC | 2,436,463,757 | 1,787 | 63,865,217 | https://www.dnazoo.org/assemblies/Canis_lupus_dingo |
| Basenji_breed-1.1 | Basenji | Luka | PacBio | 2,410,429,933 | 2,243 | 61,087,166 | GCA_004886185.2 |
| canFamDis_HiC | Golden Retriever | Scarlet | Illumina+HiC | 2,507,649,681 | 476,756 | 58,988,005 | https://www.dnazoo.org/assemblies/Canis_lupus_familiaris |

Table S4 ISO-seq reads mapping in different canine assemblies

| Assembly | BUSCO score | ISO-seq data | | | |
|---|---|---|---|---|---|
| | | N. of mapped reads (change from canFam3.1) | Mapped bases (change from canFam3.1) | N. of reads with >5% mapped based in new assembly (%) | N. of reads with >5% mapped based in canFam3.1 (%) |
| canFam3.1 | 95.20% | 532,845 | 1,533,421,214 bp | - | - |
| CanFam GSD1.0 | 95.40% | 532,959 (+0.02%) | 1,543,939,818 bp (+0.69%) | 25,609 (4.80%) | 875 (0.16%) |
| ASM864105v3 | 93.60% | 532,949 (+0.02%) | 1,543,851,433 bp (+0.68%) | 25,408 (4.76%) | 4,549 (0.85%) |
| UMICH_Zoey_3.1 | 95.20% | 532,886 (+0.01%) | 1,542,024,196 bp (+0.56%) | 25,361 (4.75%) | 2,087 (0.39%) |
| ASM325472v1_HiC | 96.00% | 532,616 (-0.04%) | 1,539,283,581 bp (+0.38%) | 25,394 (4.76%) | 7,073 (1.33%) |
| Basenji_breed-1.1 | 94.80% | 532,322 (-0.1%) | 1,523,786,336 bp (-0.63%) | 25,134 (4.71%) | 19,470 (3.65%) |
| canFamDis_HiC | 95.20% | 532,930 (+0.02%) | 1,528,715,704 bp (-0.31%) | 19,988 (3.74%) | 23,620 (4.43%) |

Table S5. Filled gaps within the cancer genes from COSMIC

| | COSMIC tier1 genes | COSMIC tier2 genes |
|---|---|---|
| Number of genes in GSD1.0 | 532 | 128 |
| Number of genes with filled gaps | 286 | 80 |
| Number of the filled gaps | 531 | 182 |
| Exons (count/length) | 328/92.2 Kb | 94/30.4 Kb |
| coding sequences (count/length) | 241/46.2 Kb | 77/18.0 Kb |
| 5'UTR (count/length) | 310/61.5 Kb | 115/21.0 Kb |
| 3'UTR (count/length) | 54/17.7 Kb | 14/2.8 Kb |

Table S6 10x sequencing of Mischka and 27 dogs

| Dog id | Breed | Sex | Access number | Sequencing depth | Average molecule length |
|---|---|---|---|---|---|
| Mischka | German Shepherd | female | SRR10428535 | 93.50X | 32515 bp |
| BOX_1 | Boxer | male | SRR10441650 | 48.19X | 49912 bp |
| CS_1 | Cocker Spaniel | female | SRR10441644 | 48.43X | 58612 bp |
| FCR_2 | Flat-coated Retriever | male | SRR10441645 | 49.31X | 51482 bp |
| SCH_1 | Schnauzer | male | SRR10441646 | 35.80X | 50857 bp |
| BOX_2 | Boxer | male | SRR10441649 | 32.15X | 51860 bp |
| FCR_1 | Flat-coated Retriever | female | SRR10441648 | 34.45X | 56222 bp |
| GR_1 | Golden Retriever | male | SRR10441647 | 45.82X | 37071 bp |
| BM_1 | Bernese Mountain | male | SRR10441652 | 36.61X | 48888 bp |
| CKCS_1 | Cavalier King Charles Spaniel | male | SRR10441637 | 30.36X | 52694 bp |
| CS_2 | Cocker Spaniel | female | SRR10441651 | 46.66X | 81696 bp |
| CS_3 | Cocker Spaniel | female | SRR10441643 | 55.82X | 50670 bp |
| DH_1 | Dachshund | female | SRR10441642 | 31.07X | 45266 bp |
| DBM_1 | Dobermann | female | SRR10441641 | 87.66X | 78645 bp |
| GSD_1 | German Shepherd | male | SRR10441640 | 58.99X | 56919 bp |
| GSD_2 | German Shepherd | female | SRR10441638 | 38.61X | 63405 bp |
| GSD_3 | German Shepherd | male | SRR10441639 | 36.95X | 72428 bp |
| GD_1 | Great Dane | female | SRR10441636 | 42.57X | 69816 bp |
| IWH_1 | Irish Wolfhound | female | SRR10441635 | 37.87X | 63311 bp |
| IWH_2 | Irish Wolfhound | female | SRR10441634 | 40.90X | 63563 bp |
| LEO_1 | Leonberger | male | SRR10441633 | 30.67X | 78586 bp |
| PUG_1 | Pug | male | SRR10441632 | 30.10X | 76245 bp |
| RTW_1 | Rottweiler | male | SRR10441631 | 32.18X | 52249 bp |
| RTW_2 | Rottweiler | male | SRR10441630 | 56.00X | 44930 bp |
| SS_1 | Springer Spaniel | female | SRR10441629 | 43.27X | 61677 bp |
| SBT_1 | Staffordshire Bull Terrier | male | SRR10441628 | 33.85X | 69289 bp |
| SP_1 | Standard Poodle | female | SRR10441627 | 43.66X | 66798 bp |
| WM_1 | Weimaraner | female | SRR10441626 | 50.76X | 59408 bp |

Table S7. Primers used for validation and genotyping

| Primer Set | Gene | ID | Sequence 5'->3' |
|---|---|---|---|
| Target Gene Expression | *CLK3* | CLK3_ex3-4-F | CGGAGATCTCGGTCCAGAAG |
| | | CLK3_ex3-4-R | CACCGGTACGTGTCACTGTC |
| | *CPLX3* | CPLX3_ex3-4-F | GATAAATATCGGCTGCCCAAG |
| | | CPLX3_ex3-4-R | CCTCCTCTGTGTCCTCCTCA |
| | *CSK* | CSK_ex7-8-F | CGTGATGCTAGGCGATTACC |
| | | CSK_ex7-8-R | AGGTTGCTATGCCGAAGTTG |
| | *CYP1A2* | CYP1A2_ex2-F | TGCCAGCAGGTGTGAGAGTA |
| | | CYP1A2_ex2-R | TGCCTTCACTTGATGGAGAA |
| | *EDC3* | EDC3_ex3-F | TGCAGAACTCCTTGCTCACA |
| | | EDC3_ex3-R | GGTACCGCCATGATGAGAAC |
| | *IGSF6* | IGSF6_ex1-2-F | GATCGTTCTTGGCCTGGA |
| | | IGSF6_ex1-2-R | ACTGAATGGTTGCAGCCTCT |
| | *MANEA* | MANEA_ex6-F | ACAGACCACCAGAGAATCTACCA |
| | | MANEA_ex6-R | GGCACTGTAGAGTTCAGGAACAG |
| | *METTL9* | METTL9_ex2-3-F | GGAGCCTCTTCCTAGCAACC |
| | | METTL9_ex2-3-R | CCTTGACCAAGGAACACAGAT |
| | *OTOA* | OTOA_ex3-4-F | CCGAATCTCTCCCAAGTCAA |
| | | OTOA_ex3-4-R | CAGATGCTCAGCAATTCCAC |
| | *POLI* | POLI_ex8-F | TCAGGAAGTGAGCGGAGTG |
| | | POLI_ex8-R | GGGAGGCTCCACGAGATACT |
| | *PPHLN1* | PPHLN1_ex9-10-F | CAATGGTGCTGTGGATCCTG |
| | | PPHLN1_ex9-10-R | GAGGCTGCAAGCAAGTGG |
| | *PRICKLE1* | PRICKLE1_ex6-7-F | TGCTGCCTTGAGTGTGAAAC |
| | | PRICKLE1_ex6-7-R | GTCATCTGGGCATGGTCAAC |
| | *RAB32* | RAB32_ex3.2-F | ATTCTTGCCAACCACCAAAG |
| | | RAB32_ex3.2-R | TCAGGGTCTCCTGATCCAGT |
| | *SCAMP2* | SCAMP2_ex3-4-F | TTCCTGTCCAGCTCTTCCTG |
| | | SCAMP2_ex3-4-R | CGACAACAGTTCCTGTCACG |
| | *ULK3* | ULK3_ex6-7-F | TCCTGGTCCTTCTTCACAGC |
| | | ULK3_ex6-7-R | CGCATCTCCTTCCAGGACT |
| | *YAF2* | YAF2_ex2-3-F | CCCTACACAGTCAAAGAAAGAGAAA |
| | | YAF2_ex2-3-R | AGCACTACTCCGATCCACATTT |
| | *ZCRB1* | ZCRB1_ex5-6-F | GACAATGGAAGAGCAGCTGAG |
| | | ZCRB1_ex5-6-R | TGGAGGTTCTCGTTCTCCAA |
| Housekeeper Gene Expression | *RPS19* | RPS19-F | CTTGGAGCCTCTGCTGAAGT |
| | | RPS19-R | GGTCGGCTCCATGACTAAGA |
| | *RPS5* | RPS5-F | TCATGAGCTTCTTGCCATTG |
| | | RPS5-R | CGCTTCCGTAAGGCACAGT |
| PCR Genotyping Primers | *RAB32* | RAB32-F | GGGATGAGGCACAGAAATC |
| | | RAB32-R2 | ATGTGGTGCTCAATCTCACC |
| | *MANEA* | MANEA-F | TGTAACACAAAGTGTGTCTTTAACT |
| | | MANEA-R2 | TCTCTGGTGGTCTGTTAGAGC |
| | *POLI* | POLI-R2 | CCTGAAGATAAGGAGAGCCTCA |
| | | POLI-F2 | AGCAAGAGCAGGAAGAGCAG |
| | | POLI-W1 | CCAGATTGCCAGCTCTCTT |
| | *CYP1A2* | CYP1A2-F-4 | GATGTTCCTCCCTAGCGACA |

| Primer Set | Gene | ID | Sequence 5'->3' |
|---|---|---|---|
| ddPCR Genotyping Target | | CYP1A2-R-4 | CCATGTTCTCTTGGCTCCAT |
| | | CYP1A2-probe-4 | 6FAM-TCCCTGAAGACCTGCCCTGGGT-MGBNFQ |
| | *PPHLN1* | PPHLN1-F | TCCTAATGGCCTCAGTCAGA |
| | | PPHLN1-R | CACGCAGTCTGTGGAAACA |
| | | PPHLN1-probe | 6FAM-CCGTGATTTGGACGGCTTGC-MGBNFQ |
| | *OTOA* | OTOA-F | CCTGGAATCTGGAACGCTAT |
| | | OTOA-R | ACAGGATGATGTCCCAGGAA |
| | | OTOA-Probe | 6FAM-GACTTGCCACTCCATGGCTCAGA-MGBNFQ |
| ddPCR Genotyping Reference | C7orf28B | C7orf28B-F | CAACACAGGTTGACCAAGGA |
| | | C7orf28B-R | TTGTGCAGGATCAGAGCATC |
| | | C7orf28B-probe | VIC-TGCCATTTGTGTGCATCCCCA-TAMRA |
| | RPP30 | RPP30-F | GGTCCTGGGATTTCAGCAT |
| | | RPP30-R | AGCTGCGTTCTCCACCAG |
| | | RPP30-probe | VIC-TTAGGTCGGAACCCGCTCGC-TAMRA |
| Break Point Primers | *OTOA* | OTOA-Break-F | TCCATCAAAGCCACAATGAA |
| | | OTOA-Break-R | CAAGGCCACACAGCTAGTCA |

Table S8 Gene expression summary for structural variant loci.

| SV | Locus | Location[2] | Gene | Expression p-value (samples)[1] | | |
|---|---|---|---|---|---|---|
| | | | | Liver | Spleen | Heart |
| Deletion | MANEA | chr12:54,394,997-54,440,487 | *MANEA* | 0.31 (het=6, del=14) | 0.95 (het=5, del=10) | na |
| Deletion | RAB32 | chr1:37,945,215-37,966,816 | *RAB32* | 0.17 (wt=6, del=3) | na (wt=6, del=2) | na |
| Deletion | POLI | chr1:21,276,172-21,301,190 | *POLI* | na | na | 0.95 (wt=3,het=3,del=3) |
| CNV | OTOA | chr6:23,345,645-23,406,849 | *OTOA* | na (CNV2=5; CNV>2=2) | na (CNV2=5; CNV>2=1) | na |
| | | | *IGSF6* | na (CNV2=5; CNV>2=2) | na (CNV2=5; CNV>2=1) | na |
| | | | *METTL9* | na (CNV2=5; CNV>2=2) | na (CNV2=5; CNV>2=1) | na |
| CNV | PPHLN1 | chr27:34,959,082-35,101,842 | *PPHLN1* | 0.64 (CNV2=13; CNV>2=7) | 0.54 (CNV2=8; CNV>2=7) | na |
| | | | *YAF* | 0.49 (CNV2=13; CNV>2=7) | 0.73 (CNV2=4; CNV>2=4) | na |
| | | | *ZCRB1* | 0.13 (CNV2=13; CNV>2=7) | 0.614 (CNV2=8; CNV>2=7) | na |
| CNV | CYP1A2 | chr30:38,258,389-38,264,108 | *CYP1A2* | 0.66 (CNV3=7; CNV>3=11) | 0.90 (CNV3=5; CNV>3=9) | na |
| | | | *CLK3* | 0.28 (CNV3=7; CNV>3=8) | 0.28 (CNV3=5; CNV>3=8) | na |
| | | | *CSK* | 0.54 (CNV3=7; CNV>3=9) | 0.61 (CNV3=5; CNV>3=9) | na |
| | | | *SCAMP2* | 0.45 (CNV3=6; CNV>3=7) | 0.11 (CNV3=4; CNV>3=4) | na |
| | | | *ULK3* | na (CNV3=2; CNV>3=2) | 0.11 (CNV3=2; CNV>3=6) | na |

1Samples available per locus per genotype; SV, structural variant type; na, not applicable. 2Location with GSD1.0 co-ordinates. For PPHLN1 and CYP1A2, the break point was determined from reads, not PCR validation

Table S9 Sequencing data generated from three HiC libraries for Mischka dog

| library | Number of read pairs | Bases count (bp) | Coverage | Accession |
|---|---|---|---|---|
| hic1 | 118,370,222 | 35,747,807,044 | 14.30X | SRR10428538 |
| hic2 | 133,709,087 | 40,380,144,274 | 16.15X | SRR10428537 |
| hic3 | 150,143,152 | 45,343,231,904 | 18.14X | SRR10428536 |

Table S10 Tissues used for nanopore sequencing

| Tissue | female RIN | male RIN |
|---|---|---|
| heart | 8 | 7,1 |
| lung | 5,2 | 7,7 |
| liver | 3,4 | 5 |
| spleen | 4,8 | 4,2 |
| pancreas | 3,9 | 6,2 |
| kidney | 3,5 | 3,2 |
| skeletal muscles | 7,8 | 6 |
| testis and ovary | 5,8 | 5,8 |
| occipital cortex | 7,6 | 6,3 |
| parietal lobe | 8,2 | 7,7 |
| frontal lobe | 8,6 | 9,1 |
| temporal cortex | 7,9 | 8 |
| thalamus | 7,9 | 5,8 |
| hypothalamus | 8,8 | 7,7 |
| superior colliculus | 8,1 | 8 |
| olfactory bulb | 8,7 | 9,1 |
| pineal | 8,4 | 7,2 |
| cerebellum | 8,2 | 6,9 |
| midbrain | 7,9 | 6,6 |
| pons | 8,5 | 8 |
| medulla oblongata | 7,9 | 7,6 |
| spinal cord | 8,6 | 7,2 |
| piriform cortex | 8,7 | 8,5 |

### Table S11 Datasets used for annotation

| Accession | Data_type | Sample information | Instrument model | Bases count (Mb) |
|---|---|---|---|---|
| SRR5889350 | RNA-seq | RNA-Seq: domestic dog - adult stomach | Illumina HiSeq 2500 | 116051 |
| SRR5889349 | RNA-seq | RNA-Seq: domestic dog - adult thyroid | Illumina HiSeq 2500 | 86340 |
| SRR5889348 | RNA-seq | RNA-Seq: domestic dog - adult small intenstine | Illumina HiSeq 2500 | 89337 |
| SRR5889347 | RNA-seq | RNA-Seq: domestic dog - adult spleen | Illumina HiSeq 2500 | 75150 |
| SRR5889346 | RNA-seq | RNA-Seq: domestic dog - adult skeletal muscle | Illumina HiSeq 2500 | 78611 |
| SRR5889345 | RNA-seq | RNA-Seq: domestic dog - adult skin | Illumina HiSeq 2500 | 79376 |
| SRR5889344 | RNA-seq | RNA-Seq: domestic dog - adult salivary gland | Illumina HiSeq 2500 | 85933 |
| SRR5889343 | RNA-seq | RNA-Seq: domestic dog - adult right ventricle | Illumina HiSeq 2500 | 79852 |
| SRR5889342 | RNA-seq | RNA-Seq: domestic dog - adult right atrium | Illumina HiSeq 2500 | 80202 |
| SRR5889341 | RNA-seq | RNA-Seq: domestic dog - adult pituitary gland | Illumina HiSeq 2500 | 76105 |
| SRR5889340 | RNA-seq | RNA-Seq: domestic dog - adult pancreas | Illumina HiSeq 2500 | 79995 |
| SRR5889339 | RNA-seq | RNA-Seq: domestic dog - adult lymph node | Illumina HiSeq 2500 | 89339 |
| SRR5889338 | RNA-seq | RNA-Seq: domestic dog - adult lung | Illumina HiSeq 2500 | 63987 |
| SRR5889337 | RNA-seq | RNA-Seq: domestic dog - adult liver | Illumina HiSeq 2500 | 74845 |
| SRR5889336 | RNA-seq | RNA-Seq: domestic dog - adult left ventricle | Illumina HiSeq 2500 | 73207 |
| SRR5889335 | RNA-seq | RNA-Seq: domestic dog - adult left atrium | Illumina HiSeq 2500 | 78578 |
| SRR5889334 | RNA-seq | RNA-Seq: domestic dog - dev. day 44 liver | Illumina HiSeq 2500 | 171683 |
| SRR5889333 | RNA-seq | RNA-Seq: domestic dog - dev. day 44 lung | Illumina HiSeq 2500 | 164518 |
| SRR5889332 | RNA-seq | RNA-Seq: domestic dog - dev. day 44 heart | Illumina HiSeq 2500 | 191463 |
| SRR5889331 | RNA-seq | RNA-Seq: domestic dog - dev. day 44 kidney | Illumina HiSeq 2500 | 176577 |
| SRR5889330 | RNA-seq | RNA-Seq: domestic dog - dev. day 39 liver | Illumina HiSeq 2500 | 164358 |
| SRR5889329 | RNA-seq | RNA-Seq: domestic dog - dev. day 39 head | Illumina HiSeq 2500 | 226303 |
| SRR5889328 | RNA-seq | RNA-Seq: domestic dog - dev. day 39 kidney | Illumina HiSeq 2500 | 154766 |
| SRR5889327 | RNA-seq | RNA-Seq: domestic dog - dev. day 39 lung | Illumina HiSeq 2500 | 162132 |
| SRR5889326 | RNA-seq | RNA-Seq: domestic dog - adult adipose | Illumina HiSeq 2500 | 79339 |
| SRR5889325 | RNA-seq | RNA-Seq: domestic dog - dev. day 39 heart | Illumina HiSeq 2500 | 69449 |
| SRR5889324 | RNA-seq | RNA-Seq: domestic dog - dev. day 33 head | Illumina HiSeq 2500 | 154182 |
| SRR5889323 | RNA-seq | RNA-Seq: domestic dog - dev. day 33 lung | Illumina HiSeq 2500 | 164710 |
| SRR5889322 | RNA-seq | RNA-Seq: domestic dog - dev. day 36 liver | Illumina HiSeq 2500 | 166856 |
| SRR5889321 | RNA-seq | RNA-Seq: domestic dog - dev. day 44 head | Illumina HiSeq 2500 | 159356 |
| SRR5889320 | RNA-seq | RNA-Seq: domestic dog - dev. day 36 kidney | Illumina HiSeq 2500 | 166900 |
| SRR5889319 | RNA-seq | RNA-Seq: domestic dog - dev. day 36 heart | Illumina HiSeq 2500 | 171619 |
| SRR5889318 | RNA-seq | RNA-Seq: domestic dog - dev. day 36 lung | Illumina HiSeq 2500 | 151263 |
| SRR5889317 | RNA-seq | RNA-Seq: domestic dog - dev. day 36 head | Illumina HiSeq 2500 | 156939 |
| SRR5889316 | RNA-seq | RNA-Seq: domestic dog - dev. day 33 heart | Illumina HiSeq 2500 | 139957 |
| SRR5889315 | RNA-seq | RNA-Seq: domestic dog - dev. day 33 liver | Illumina HiSeq 2500 | 159043 |
| SRR5889314 | RNA-seq | RNA-Seq: domestic dog - adult kidney cortex | Illumina HiSeq 2500 | 78551 |
| SRR5889313 | RNA-seq | RNA-Seq: domestic dog - adult kidney medulla | Illumina HiSeq 2500 | 80414 |
| SRR5889312 | RNA-seq | RNA-Seq: domestic dog - adult adrenal gland | Illumina HiSeq 2500 | 71399 |
| SRR5889311 | RNA-seq | RNA-Seq: domestic dog - adult bladder | Illumina HiSeq 2500 | 87037 |
| SRR5889310 | RNA-seq | RNA-Seq: domestic dog - adult bone marrow | Illumina HiSeq 2500 | 92006 |
| SRR5889309 | RNA-seq | RNA-Seq: domestic dog - adult cartilage | Illumina HiSeq 2500 | 79482 |
| SRR5889308 | RNA-seq | RNA-Seq: domestic dog - adult cerebellum | Illumina HiSeq 2500 | 80283 |
| SRR5889307 | RNA-seq | RNA-Seq: domestic dog - adult colon | Illumina HiSeq 2500 | 72415 |
| SRR5889306 | RNA-seq | RNA-Seq: domestic dog - adult cortex occipital | Illumina HiSeq 2500 | 74877 |
| SRR5889305 | RNA-seq | RNA-Seq: domestic dog - adult frontal cortex | Illumina HiSeq 2500 | 119783 |
| SRR8997056 | RNA-seq | RNA-Seq: domestic dog - adult small intestine | Illumina HiSeq 2500 | 163546 |
| SRR8997055 | RNA-seq | RNA-Seq: domestic dog - adult skin | Illumina HiSeq 2500 | 165478 |
| SRR8997054 | RNA-seq | RNA-Seq: domestic dog - adult skeletal muscle | Illumina HiSeq 2500 | 220756 |
| SRR8997053 | RNA-seq | RNA-Seq: domestic dog - adult salivary gland | Illumina HiSeq 2500 | 220013 |
| SRR8997052 | RNA-seq | RNA-Seq: domestic dog - adult Adipose Tissue | Illumina HiSeq 2500 | 109400 |
| SRR8997051 | RNA-seq | RNA-Seq: domestic dog - adult thyroid gland | Illumina HiSeq 2500 | 203467 |
| SRR8997050 | RNA-seq | RNA-Seq: domestic dog - adult stomach | Illumina HiSeq 2500 | 364789 |
| SRR8997049 | RNA-seq | RNA-Seq: domestic dog - adult spleen | Illumina HiSeq 2500 | 216838 |
| SRR8997048 | RNA-seq | RNA-Seq: domestic dog - adult Bladder | Illumina HiSeq 2500 | 105309 |
| SRR8997047 | RNA-seq | RNA-Seq: domestic dog - adult Adrenal Gland | Illumina HiSeq 2500 | 95315 |
| SRR8997046 | RNA-seq | RNA-Seq: domestic dog - adult thyroid gland | Illumina HiSeq 2500 | 97815 |
| SRR8997045 | RNA-seq | RNA-Seq: domestic dog - adult stomach | Illumina HiSeq 2500 | 112147 |

| Accession | Data_type | Sample information | Instrument model | Bases count (Mb) |
|---|---|---|---|---|
| SRR8997044 | RNA-seq | RNA-Seq: domestic dog - adult spleen | Illumina HiSeq 2500 | 54439 |
| SRR8997043 | RNA-seq | RNA-Seq: domestic dog - adult skin | Illumina HiSeq 2500 | 87265 |
| SRR8997042 | RNA-seq | RNA-Seq: domestic dog - adult Liver | Illumina HiSeq 2500 | 104158 |
| SRR8997041 | RNA-seq | RNA-Seq: domestic dog - adult Lung | Illumina HiSeq 2500 | 128615 |
| SRR8997040 | RNA-seq | RNA-Seq: domestic dog - adult Kidney Cortex | Illumina HiSeq 2500 | 108884 |
| SRR8997039 | RNA-seq | RNA-Seq: domestic dog - adult Kidney Medulla | Illumina HiSeq 2500 | 98333 |
| SRR8997038 | RNA-seq | RNA-Seq: domestic dog - adult Left Atrium | Illumina HiSeq 2500 | 103694 |
| SRR8997037 | RNA-seq | RNA-Seq: domestic dog - adult Left Ventricle | Illumina HiSeq 2500 | 80631 |
| SRR8997036 | RNA-seq | RNA-Seq: domestic dog - adult Bone Marrow | Illumina HiSeq 2500 | 106469 |
| SRR8997035 | RNA-seq | RNA-Seq: domestic dog - adult Cerebellum | Illumina HiSeq 2500 | 104139 |
| SRR8997034 | RNA-seq | RNA-Seq: domestic dog - adult Colon | Illumina HiSeq 2500 | 105988 |
| SRR8997033 | RNA-seq | RNA-Seq: domestic dog - adult Frontal Cortex | Illumina HiSeq 2500 | 95478 |
| SRR8997032 | RNA-seq | RNA-Seq: domestic dog - adult Spleen | Illumina HiSeq 2500 | 95278 |
| SRR8997031 | RNA-seq | RNA-Seq: domestic dog - adult Small Intestine | Illumina HiSeq 2500 | 88758 |
| SRR8997030 | RNA-seq | RNA-Seq: domestic dog - adult Occipital Lobe | Illumina HiSeq 2500 | 102884 |
| SRR8997029 | RNA-seq | RNA-Seq: domestic dog - adult Lymph node | Illumina HiSeq 2500 | 86460 |
| SRR8997028 | RNA-seq | RNA-Seq: domestic dog - adult Right Atrium | Illumina HiSeq 2500 | 80039 |
| SRR8997027 | RNA-seq | RNA-Seq: domestic dog - adult Pancreas | Illumina HiSeq 2500 | 120908 |
| SRR8997026 | RNA-seq | RNA-Seq: domestic dog - adult Salivary Gland | Illumina HiSeq 2500 | 106087 |
| SRR8997025 | RNA-seq | RNA-Seq: domestic dog - adult Right Ventricle | Illumina HiSeq 2500 | 100118 |
| SRR8997024 | RNA-seq | RNA-Seq: domestic dog - adult Skin | Illumina HiSeq 2500 | 103969 |
| SRR8997023 | RNA-seq | RNA-Seq: domestic dog - adult Skeletal Muscle | Illumina HiSeq 2500 | 113644 |
| SRR8997022 | RNA-seq | RNA-Seq: domestic dog - adult colon | Illumina HiSeq 2500 | 69816 |
| SRR8997021 | RNA-seq | RNA-Seq: domestic dog - adult frontal cortex | Illumina HiSeq 2500 | 68577 |
| SRR8997020 | RNA-seq | RNA-Seq: domestic dog - adult adipose | Illumina HiSeq 2500 | 37489 |
| SRR8997019 | RNA-seq | RNA-Seq: domestic dog - adult adrenal gland | Illumina HiSeq 2500 | 56900 |
| SRR8997018 | RNA-seq | RNA-Seq: domestic dog - adult Stomach | Illumina HiSeq 2500 | 116943 |
| SRR8997017 | RNA-seq | RNA-Seq: domestic dog - adult Thyroid | Illumina HiSeq 2500 | 101656 |
| SRR8997016 | RNA-seq | RNA-Seq: domestic dog - adult cartilage | Illumina HiSeq 2500 | 59847 |
| SRR8997015 | RNA-seq | RNA-Seq: domestic dog - adult cerebellum | Illumina HiSeq 2500 | 74264 |
| SRR8997014 | RNA-seq | RNA-Seq: domestic dog - adult bladder | Illumina HiSeq 2500 | 65516 |
| SRR8997013 | RNA-seq | RNA-Seq: domestic dog - adult bone marrow | Illumina HiSeq 2500 | 59299 |
| SRR8997012 | RNA-seq | RNA-Seq: domestic dog - adult occipital cortex | Illumina HiSeq 2500 | 80151 |
| SRR8997011 | RNA-seq | RNA-Seq: domestic dog - adult lymph node | Illumina HiSeq 2500 | 62963 |
| SRR8997010 | RNA-seq | RNA-Seq: domestic dog - adult lung | Illumina HiSeq 2500 | 69661 |
| SRR8997009 | RNA-seq | RNA-Seq: domestic dog - adult liver | Illumina HiSeq 2500 | 79360 |
| SRR8997008 | RNA-seq | RNA-Seq: domestic dog - adult left ventricle | Illumina HiSeq 2500 | 65217 |
| SRR8997007 | RNA-seq | RNA-Seq: domestic dog - adult left atrium | Illumina HiSeq 2500 | 70367 |
| SRR8997006 | RNA-seq | RNA-Seq: domestic dog - adult kidney medulla | Illumina HiSeq 2500 | 60191 |
| SRR8997005 | RNA-seq | RNA-Seq: domestic dog - adult kidney cortex | Illumina HiSeq 2500 | 70913 |
| SRR8997004 | RNA-seq | RNA-Seq: domestic dog - adult pituitary gland | Illumina HiSeq 2500 | 53180 |
| SRR8997003 | RNA-seq | RNA-Seq: domestic dog - adult pancreas | Illumina HiSeq 2500 | 64993 |
| SRR8997002 | RNA-seq | RNA-Seq: domestic dog - adult kidney medulla | Illumina HiSeq 2500 | 89056 |
| SRR8997001 | RNA-seq | RNA-Seq: domestic dog - adult kidney cortex | Illumina HiSeq 2500 | 116731 |
| SRR8997000 | RNA-seq | RNA-Seq: domestic dog - adult colon | Illumina HiSeq 2500 | 90337 |
| SRR8996999 | RNA-seq | RNA-Seq: domestic dog - adult cerebellum | Illumina HiSeq 2500 | 84790 |
| SRR8996998 | RNA-seq | RNA-Seq: domestic dog - adult cortex occipital | Illumina HiSeq 2500 | 83610 |
| SRR8996997 | RNA-seq | RNA-Seq: domestic dog - adult cortex frontal | Illumina HiSeq 2500 | 99968 |
| SRR8996996 | RNA-seq | RNA-Seq: domestic dog - adult bladder | Illumina HiSeq 2500 | 51771 |
| SRR8996995 | RNA-seq | RNA-Seq: domestic dog - adult adrenal gland | Illumina HiSeq 2500 | 113118 |
| SRR8996994 | RNA-seq | RNA-Seq: domestic dog - adult cartilage | Illumina HiSeq 2500 | 108910 |
| SRR8996993 | RNA-seq | RNA-Seq: domestic dog - adult bone marrow | Illumina HiSeq 2500 | 145990 |
| SRR8996992 | RNA-seq | RNA-Seq: domestic dog - adult right atrium | Illumina HiSeq 2500 | 60862 |
| SRR8996991 | RNA-seq | RNA-Seq: domestic dog - adult right ventricle | Illumina HiSeq 2500 | 78180 |
| SRR8996990 | RNA-seq | RNA-Seq: domestic dog - adult salivary gland | Illumina HiSeq 2500 | 66129 |
| SRR8996989 | RNA-seq | RNA-Seq: domestic dog - adult skeletal muscle | Illumina HiSeq 2500 | 43949 |
| SRR8996988 | RNA-seq | RNA-Seq: domestic dog - adult skin | Illumina HiSeq 2500 | 61603 |
| SRR8996987 | RNA-seq | RNA-Seq: domestic dog - adult small intestine | Illumina HiSeq 2500 | 55038 |
| SRR8996986 | RNA-seq | RNA-Seq: domestic dog - adult spleen | Illumina HiSeq 2500 | 68625 |
| SRR8996985 | RNA-seq | RNA-Seq: domestic dog - adult stomach | Illumina HiSeq 2500 | 68866 |

| Accession | Data_type | Sample information | Instrument model | Bases count (Mb) |
|---|---|---|---|---|
| SRR8996984 | RNA-seq | RNA-Seq: domestic dog - adult thyroid gland | Illumina HiSeq 2500 | 56031 |
| SRR8996983 | RNA-seq | RNA-Seq: domestic dog - adult adipose | Illumina HiSeq 2500 | 91208 |
| SRR8996982 | RNA-seq | RNA-Seq: domestic dog - adult pancreas | Illumina HiSeq 2500 | 201622 |
| SRR8996981 | RNA-seq | RNA-Seq: domestic dog - adult pituitary gland | Illumina HiSeq 2500 | 153350 |
| SRR8996980 | RNA-seq | RNA-Seq: domestic dog - adult lung | Illumina HiSeq 2500 | 277078 |
| SRR8996979 | RNA-seq | RNA-Seq: domestic dog - adult lymph node | Illumina HiSeq 2500 | 208554 |
| SRR8996978 | RNA-seq | RNA-Seq: domestic dog - adult left ventricle | Illumina HiSeq 2500 | 142417 |
| SRR8996977 | RNA-seq | RNA-Seq: domestic dog - adult liver | Illumina HiSeq 2500 | 209869 |
| SRR8996976 | RNA-seq | RNA-Seq: domestic dog - adult kidney medulla | Illumina HiSeq 2500 | 152046 |
| SRR8996975 | RNA-seq | RNA-Seq: domestic dog - adult left atrium | Illumina HiSeq 2500 | 195671 |
| SRR8996974 | RNA-seq | RNA-Seq: domestic dog - adult right atrium | Illumina HiSeq 2500 | 180801 |
| SRR8996973 | RNA-seq | RNA-Seq: domestic dog - adult right ventricle | Illumina HiSeq 2500 | 165186 |
| SRR8996972 | RNA-seq | RNA-Seq: domestic dog - adult salivary gland | Illumina HiSeq 2500 | 87378 |
| SRR8996971 | RNA-seq | RNA-Seq: domestic dog - adult skeletal muscle | Illumina HiSeq 2500 | 92176 |
| SRR8996970 | RNA-seq | RNA-Seq: domestic dog - adult pituitary gland | Illumina HiSeq 2500 | 98085 |
| SRR8996969 | RNA-seq | RNA-Seq: domestic dog - adult right ventricle | Illumina HiSeq 2500 | 94230 |
| SRR8996968 | RNA-seq | RNA-Seq: domestic dog - adult lymph node | Illumina HiSeq 2500 | 126212 |
| SRR8996967 | RNA-seq | RNA-Seq: domestic dog - adult pancreas | Illumina HiSeq 2500 | 111672 |
| SRR8996966 | RNA-seq | RNA-Seq: domestic dog - adult liver | Illumina HiSeq 2500 | 104418 |
| SRR8996965 | RNA-seq | RNA-Seq: domestic dog - adult lung | Illumina HiSeq 2500 | 93260 |
| SRR8996964 | RNA-seq | RNA-Seq: domestic dog - adult left atrium | Illumina HiSeq 2500 | 94070 |
| SRR8996963 | RNA-seq | RNA-Seq: domestic dog - adult left ventricle | Illumina HiSeq 2500 | 92902 |
| SRR8996962 | RNA-seq | RNA-Seq: domestic dog - adult cartilage | Illumina HiSeq 2500 | 179328 |
| SRR8996961 | RNA-seq | RNA-Seq: domestic dog - adult cerebellum | Illumina HiSeq 2500 | 169980 |
| SRR8996960 | RNA-seq | RNA-Seq: domestic dog - adult colon | Illumina HiSeq 2500 | 207710 |
| SRR8996959 | RNA-seq | RNA-Seq: domestic dog - adult cortex frontal | Illumina HiSeq 2500 | 211637 |
| SRR8996958 | RNA-seq | RNA-Seq: domestic dog - adult adipose | Illumina HiSeq 2500 | 206430 |
| SRR8996957 | RNA-seq | RNA-Seq: domestic dog - adult adrenal gland | Illumina HiSeq 2500 | 203450 |
| SRR8996956 | RNA-seq | RNA-Seq: domestic dog - adult bladder | Illumina HiSeq 2500 | 166225 |
| SRR8996955 | RNA-seq | RNA-Seq: domestic dog - adult bone marrow | Illumina HiSeq 2500 | 220237 |
| SRR8996954 | RNA-seq | RNA-Seq: domestic dog - adult cortex occipital | Illumina HiSeq 2500 | 233810 |
| SRR8996953 | RNA-seq | RNA-Seq: domestic dog - adult kidney cortex | Illumina HiSeq 2500 | 182267 |
| SRR9035349 | ATAC-seq | ATAC-seq of adult canine pancreas | Illumina HiSeq 2500 | 26536 |
| SRR9035348 | ATAC-seq | ATAC-seq of adult canine pituitary | Illumina HiSeq 2500 | 24291 |
| SRR9035347 | ATAC-seq | ATAC-seq of adult canine spleen | Illumina HiSeq 2500 | 27958 |
| SRR9035346 | ATAC-seq | ATAC-seq of adult canine stomach | Illumina HiSeq 2500 | 14562 |
| SRR9035345 | ATAC-seq | ATAC-seq of adult canine bone_marrow | Illumina HiSeq 2500 | 27715 |
| SRR9035344 | ATAC-seq | ATAC-seq of adult canine small_intestine | Illumina HiSeq 2500 | 28104 |
| SRR9035343 | ATAC-seq | ATAC-seq of adult canine spleen | Illumina HiSeq 2500 | 19544 |
| SRR9035342 | ATAC-seq | ATAC-seq of adult canine pituitary | Illumina HiSeq 2500 | 27147 |
| SRR9035341 | ATAC-seq | ATAC-seq of adult canine salivary | Illumina HiSeq 2500 | 19779 |
| SRR9035340 | ATAC-seq | ATAC-seq of adult canine lymph_node | Illumina HiSeq 2500 | 25757 |
| SRR9035339 | ATAC-seq | ATAC-seq of adult canine pancreas | Illumina HiSeq 2500 | 30839 |
| SRR9035338 | ATAC-seq | ATAC-seq of adult canine left_atrium | Illumina HiSeq 2500 | 29319 |
| SRR9035337 | ATAC-seq | ATAC-seq of adult canine liver | Illumina HiSeq 2500 | 26550 |
| SRR9035336 | ATAC-seq | ATAC-seq of adult canine pancreas | Illumina HiSeq 2500 | 28673 |
| SRR9035335 | ATAC-seq | ATAC-seq of adult canine salivary | Illumina HiSeq 2500 | 26858 |
| SRR9035334 | ATAC-seq | ATAC-seq of adult canine liver | Illumina HiSeq 2500 | 18324 |
| SRR9035333 | ATAC-seq | ATAC-seq of adult canine lymph_node | Illumina HiSeq 2500 | 20378 |
| SRR9035332 | ATAC-seq | ATAC-seq of adult canine skeletal_muscle | Illumina HiSeq 2500 | 10224 |
| SRR9035331 | ATAC-seq | ATAC-seq of adult canine salivary | Illumina HiSeq 2500 | 31177 |
| SRR9035330 | ATAC-seq | ATAC-seq of adult canine lymph_node | Illumina HiSeq 2500 | 25415 |
| SRR9035329 | ATAC-seq | ATAC-seq of adult canine left_atrium | Illumina HiSeq 2500 | 11829 |
| SRR9035328 | ATAC-seq | ATAC-seq of adult canine thyroid | Illumina HiSeq 2500 | 26892 |
| SRR9035327 | ATAC-seq | ATAC-seq of adult canine stomach | Illumina HiSeq 2500 | 33428 |
| SRR9035326 | ATAC-seq | ATAC-seq of adult canine right_ventricle | Illumina HiSeq 2500 | 11382 |
| SRR9035325 | ATAC-seq | ATAC-seq of adult canine pituitary | Illumina HiSeq 2500 | 25919 |
| SRR9035324 | ATAC-seq | ATAC-seq of adult canine pancreas | Illumina HiSeq 2500 | 29014 |
| SRR9035323 | ATAC-seq | ATAC-seq of adult canine occipital_cortex | Illumina HiSeq 2500 | 10767 |
| SRR9035322 | ATAC-seq | ATAC-seq of adult canine liver | Illumina HiSeq 2500 | 48261 |

| Accession | Data_type | Sample information | Instrument model | Bases count (Mb) |
|---|---|---|---|---|
| SRR9035321 | ATAC-seq | ATAC-seq of adult canine lymph_node | Illumina HiSeq 2500 | 24466 |
| SRR9035320 | ATAC-seq | ATAC-seq of adult canine right_ventricle | Illumina HiSeq 2500 | 26721 |
| SRR9035319 | ATAC-seq | ATAC-seq of adult canine salivary | Illumina HiSeq 2500 | 41628 |
| SRR9035318 | ATAC-seq | ATAC-seq of adult canine spleen | Illumina HiSeq 2500 | 26864 |
| SRR9035317 | ATAC-seq | ATAC-seq of adult canine bladder | Illumina HiSeq 2500 | 26859 |
| SRR9035316 | ATAC-seq | ATAC-seq of adult canine bone_marrow | Illumina HiSeq 2500 | 26062 |
| SRR9035315 | ATAC-seq | ATAC-seq of adult canine liver | Illumina HiSeq 2500 | 34665 |
| SRR9035314 | ATAC-seq | ATAC-seq of adult canine lymph_node | Illumina HiSeq 2500 | 23814 |
| SRR9035313 | ATAC-seq | ATAC-seq of adult canine pancreas | Illumina HiSeq 2500 | 59298 |
| SRR3727705 | RNA-seq | RNA-Seq:ADRENALGLAND:Bernese_Mountain_Dog | Illumina HiSeq 2000 | 83260 |
| SRR3727706 | RNA-seq | RNA-Seq:CEREBELLUM:Belgian_Shepherd | Illumina HiSeq 2000 | 80221 |
| SRR3727707 | RNA-seq | RNA-Seq:NOSE02:Labrador | Illumina HiSeq 2000 | 137726 |
| SRR3727708 | RNA-seq | RNA-Seq:NOSE03:Labrador | Illumina HiSeq 2000 | 139771 |
| SRR3727709 | RNA-seq | RNA-Seq:OLFBULB:Great_Swiss_Mountain_Dog | Illumina HiSeq 2000 | 69615 |
| SRR3727710 | RNA-seq | RNA-Seq:PANCREAS:Belgian_Shepherd | Illumina HiSeq 2000 | 71701 |
| SRR3727711 | RNA-seq | RNA-Seq:RETINA:Border_Collie | Illumina HiSeq 2000 | 76730 |
| SRR3727713 | RNA-seq | RNA-Seq:SKIN:Beagle | Illumina HiSeq 2000 | 79459 |
| SRR3727714 | RNA-seq | RNA-Seq:SKIN:Great_Swiss_Mountain_Dog | Illumina HiSeq 2000 | 74603 |
| SRR3727715 | RNA-seq | RNA-Seq:SPINALCORD:Great_Swiss_Mountain_Dog | Illumina HiSeq 2000 | 71203 |
| SRR3727716 | RNA-seq | RNA-Seq:SPLEEN:Belgian_Shepherd | Illumina HiSeq 2000 | 75399 |
| SRR3727717 | RNA-seq | RNA-Seq:THYMUS:Saluki | Illumina HiSeq 2000 | 77640 |
| SRR3727718 | RNA-seq | RNA-Seq:CEREBELLUM:Great_Swiss_Mountain_Dog | Illumina HiSeq 2000 | 68252 |
| SRR3727719 | RNA-seq | RNA-Seq:CORTEX:Belgian_Shepherd | Illumina HiSeq 2000 | 62806 |
| SRR3727720 | RNA-seq | RNA-Seq:GUTCOLON:Bernese_Mountain_Dog | Illumina HiSeq 2000 | 79512 |
| SRR3727722 | RNA-seq | RNA-Seq:HAIRFOLLICULE:Labrador | Illumina HiSeq 2000 | 92303 |
| SRR3727723 | RNA-seq | RNA-Seq:JEJUNUM:Labrador | Illumina HiSeq 2000 | 76866 |
| SRR3727724 | RNA-seq | RNA-Seq:KERATINOCYTE:Beagle | Illumina HiSeq 2000 | 82813 |
| SRR3727725 | RNA-seq | RNA-Seq:MAMMARYGLAND:Great_Swiss_Mountain_Dog | Illumina HiSeq 2000 | 67412 |
| SRR3727726 | RNA-seq | RNA-Seq:NOSE01:Labrador | Illumina HiSeq 2000 | 118861 |
| SRR388747 | RNA-seq | RNA Sequencing of Canis familiaris (Dog) | Illumina HiSeq 2000 | 24244 |
| SRR388735 | RNA-seq | RNA Sequencing of Canis familiaris (Dog) | Illumina HiSeq 2000 | 24350 |
| SRR388734 | RNA-seq | RNA Sequencing of Canis familiaris (Dog) | Illumina HiSeq 2000 | 24294 |
| SRR388752 | RNA-seq | RNA Sequencing of Canis familiaris (Dog) | Illumina HiSeq 2000 | 21594 |
| SRR388741 | RNA-seq | RNA Sequencing of Canis familiaris (Dog) | Illumina HiSeq 2000 | 21657 |
| SRR388736 | RNA-seq | RNA Sequencing of Canis familiaris (Dog) | Illumina HiSeq 2000 | 21641 |
| SRR388766 | RNA-seq | RNA Sequencing of Canis familiaris (Dog) | Illumina HiSeq 2000 | 19413 |
| SRR388737 | RNA-seq | RNA Sequencing of Canis familiaris (Dog) | Illumina HiSeq 2000 | 19489 |
| SRR388740 | RNA-seq | RNA Sequencing of Canis familiaris (Dog) | Illumina HiSeq 2000 | 19462 |
| SRR388760 | RNA-seq | RNA Sequencing of Canis familiaris (Dog) | Illumina HiSeq 2000 | 20233 |
| SRR388751 | RNA-seq | RNA Sequencing of Canis familiaris (Dog) | Illumina HiSeq 2000 | 20211 |
| SRR388738 | RNA-seq | RNA Sequencing of Canis familiaris (Dog) | Illumina HiSeq 2000 | 20099 |
| SRR388759 | RNA-seq | RNA Sequencing of Canis familiaris (Dog) | Illumina HiSeq 2000 | 11780 |
| SRR388739 | RNA-seq | RNA Sequencing of Canis familiaris (Dog) | Illumina HiSeq 2000 | 11702 |
| SRR388748 | RNA-seq | RNA Sequencing of Canis familiaris (Dog) | Illumina HiSeq 2000 | 11438 |
| SRR388764 | RNA-seq | RNA Sequencing of Canis familiaris (Dog) | Illumina HiSeq 2000 | 21952 |
| SRR388756 | RNA-seq | RNA Sequencing of Canis familiaris (Dog) | Illumina HiSeq 2000 | 21806 |
| SRR388742 | RNA-seq | RNA Sequencing of Canis familiaris (Dog) | Illumina HiSeq 2000 | 21898 |
| SRR388761 | RNA-seq | RNA Sequencing of Canis familiaris (Dog) | Illumina HiSeq 2000 | 21451 |
| SRR388743 | RNA-seq | RNA Sequencing of Canis familiaris (Dog) | Illumina HiSeq 2000 | 21483 |
| SRR388744 | RNA-seq | RNA Sequencing of Canis familiaris (Dog) | Illumina HiSeq 2000 | 21509 |
| SRR388762 | RNA-seq | RNA Sequencing of Canis familiaris (Dog) | Illumina HiSeq 2000 | 21925 |
| SRR388750 | RNA-seq | RNA Sequencing of Canis familiaris (Dog) | Illumina HiSeq 2000 | 22001 |
| SRR388745 | RNA-seq | RNA Sequencing of Canis familiaris (Dog) | Illumina HiSeq 2000 | 21969 |
| SRR388758 | RNA-seq | RNA Sequencing of Canis familiaris (Dog) | Illumina HiSeq 2000 | 23873 |
| SRR388755 | RNA-seq | RNA Sequencing of Canis familiaris (Dog) | Illumina HiSeq 2000 | 23905 |
| SRR388746 | RNA-seq | RNA Sequencing of Canis familiaris (Dog) | Illumina HiSeq 2000 | 23949 |
| SRR388757 | RNA-seq | RNA Sequencing of Canis familiaris (Dog) | Illumina HiSeq 2000 | 23426 |
| SRR388753 | RNA-seq | RNA Sequencing of Canis familiaris (Dog) | Illumina HiSeq 2000 | 23270 |
| SRR388749 | RNA-seq | RNA Sequencing of Canis familiaris (Dog) | Illumina HiSeq 2000 | 23386 |

| Accession | Data_type | Sample information | Instrument model | Bases count (Mb) |
|---|---|---|---|---|
| SRR388765 | RNA-seq | RNA Sequencing of Canis familiaris (Dog) | Illumina HiSeq 2000 | 19870 |
| SRR388763 | RNA-seq | RNA Sequencing of Canis familiaris (Dog) | Illumina HiSeq 2000 | 19865 |
| SRR388754 | RNA-seq | RNA Sequencing of Canis familiaris (Dog) | Illumina HiSeq 2000 | 19786 |
| SRR543733 | RNA-seq | Ross Swofford_Zyagen2_Brain_1201 | Illumina HiSeq 2000 | 30549 |
| SRR536883 | RNA-seq | Ross Swofford_Zyagen2_Brain_1201 | Illumina HiSeq 2000 | 29846 |
| SRR536881 | RNA-seq | Ross Swofford_Zyagen2_Brain_1201 | Illumina HiSeq 2000 | 31582 |
| SRR543734 | RNA-seq | Ross Swofford_Zyagen2_Skin_1202 | Illumina HiSeq 2000 | 26165 |
| SRR543732 | RNA-seq | Ross Swofford_Zyagen2_Skin_1202 | Illumina HiSeq 2000 | 26798 |
| SRR536884 | RNA-seq | Ross Swofford_Zyagen2_Skin_1202 | Illumina HiSeq 2000 | 27757 |
| SRR543735 | RNA-seq | Ross Swofford_Zyagen2_Kidney_1203 | Illumina HiSeq 2000 | 33726 |
| SRR536885 | RNA-seq | Ross Swofford_Zyagen2_Kidney_1203 | Illumina HiSeq 2000 | 32709 |
| SRR536882 | RNA-seq | Ross Swofford_Zyagen2_Kidney_1203 | Illumina HiSeq 2000 | 31960 |
| SRR8474278 | RNA-seq | L16_1377_Dog_4_Spleen; Canis lupus familiaris; RNA-Seq | Illumina HiSeq 2500 | 83329 |
| SRR8474282 | RNA-seq | L16_1513_Dog_6_Colon; Canis lupus familiaris; RNA-Seq | Illumina HiSeq 2500 | 112365 |
| SRR8474283 | RNA-seq | L16_1593_Dog_6_Heart; Canis lupus familiaris; RNA-Seq | Illumina HiSeq 2500 | 134435 |
| SRR8474284 | RNA-seq | L16_1303_Dog_6_Liver; Canis lupus familiaris; RNA-Seq | Illumina HiSeq 2500 | 55722 |
| SRR8474286 | RNA-seq | L16_1390_Dog_6_Muscle; Canis lupus familiaris; RNA-Seq | Illumina HiSeq 2500 | 69030 |
| SRR8474288 | RNA-seq | L16_1610_Dog_6_Spleen; Canis lupus familiaris; RNA-Seq | Illumina HiSeq 2500 | 111066 |
| SRR8474289 | RNA-seq | L16_1598_Dog_6_Thyroid; Canis lupus familiaris; RNA-Seq | Illumina HiSeq 2500 | 135189 |
| SRR8474291 | RNA-seq | L16_1362_Dog_7_Brain; Canis lupus familiaris; RNA-Seq | Illumina HiSeq 2500 | 65317 |
| SRR8474292 | RNA-seq | L16_1310_Dog_7_Pituitary; Canis lupus familiaris; RNA-Seq | Illumina HiSeq 2500 | 55289 |
| SRR8474293 | RNA-seq | L16_1529_Dog_8_Colon; Canis lupus familiaris; RNA-Seq | Illumina HiSeq 2500 | 137612 |
| SRR8474294 | RNA-seq | L16_1324_Dog_8_Heart; Canis lupus familiaris; RNA-Seq | Illumina HiSeq 2500 | 65858 |
| SRR8474295 | RNA-seq | L16_1305_Dog_8_Liver; Canis lupus familiaris; RNA-Seq | Illumina HiSeq 2500 | 47394 |
| SRR8474296 | RNA-seq | L16_1494_Dog_8_Lung; Canis lupus familiaris; RNA-Seq | Illumina HiSeq 2500 | 99735 |
| SRR8474298 | RNA-seq | L16_1530_Dog_8_Skin; Canis lupus familiaris; RNA-Seq | Illumina HiSeq 2500 | 103807 |
| SRR8474300 | RNA-seq | L16_1493_Dog_8_Thyroid; Canis lupus familiaris; RNA-Seq | Illumina HiSeq 2500 | 115947 |
| SRR8474301 | RNA-seq | L16_1606_Dog_9_Adipose; Canis lupus familiaris; RNA-Seq | Illumina HiSeq 2500 | 128699 |
| SRR8474302 | RNA-seq | L16_1360_Dog_10_Adipose; Canis lupus familiaris; RNA-Seq | Illumina HiSeq 2500 | 53487 |
| SRR8474297 | RNA-seq | L16_1177_Dog_8_Muscle; Canis lupus familiaris; RNA-Seq | Illumina HiSeq 2500 | 235062 |
| SRR8474297 | RNA-seq | L16_1177_Dog_8_Muscle; Canis lupus familiaris; RNA-Seq | Illumina HiSeq 2500 | 235062 |
| SRR8474287 | RNA-seq | L16_1051_Dog_6_Skin; Canis lupus familiaris; RNA-Seq | Illumina HiSeq 2500 | 304271 |
| SRR5457067 | RNA-seq | RNAseq of dog granulocytes | Illumina HiSeq 2000 | 299102 |
| SRR8474290 | RNA-seq | L16_1127_Dog_4_Adrenal; Canis lupus familiaris; RNA-Seq | Illumina HiSeq 2500 | 424831 |
| SRR8474299 | RNA-seq | L16_1145_Dog_8_Spleen; Canis lupus familiaris; RNA-Seq | Illumina HiSeq 2500 | 386260 |
| SRR10488351 | RNA-seq | AIA - all cells from infected well; Canis lupus familiaris; RNA-Seq | Illumina NovaSeq 6000 | 58778 |
| SRR10488352 | RNA-seq | AIB - all cells from infected well; Canis lupus familiaris; RNA-Seq | Illumina NovaSeq 6000 | 52784 |
| SRR10488353 | RNA-seq | AIC - all cells from infected well; Canis lupus familiaris; RNA-Seq | Illumina NovaSeq 6000 | 56445 |
| SRR10488354 | RNA-seq | AID - all cells from infected well; Canis lupus familiaris; RNA-Seq | Illumina NovaSeq 6000 | 54379 |
| SRR10488355 | RNA-seq | IIA - sorted infected cells from infected well; Canis lupus familiaris; RNA-Seq | Illumina NovaSeq 6000 | 46614 |
| SRR10488356 | RNA-seq | IIB - sorted infected cells from infected well; Canis lupus familiaris; RNA-Seq | Illumina NovaSeq 6000 | 47056 |
| SRR10488357 | RNA-seq | IIC - sorted infected cells from infected well; Canis lupus familiaris; RNA-Seq | Illumina NovaSeq 6000 | 67107 |
| SRR10488358 | RNA-seq | IID - sorted infected cells from infected well; Canis lupus familiaris; RNA-Seq | Illumina NovaSeq 6000 | 48624 |
| SRR10488359 | RNA-seq | UIA - sorted uninfected cells from infected well; Canis lupus familiaris; RNA-Seq | Illumina NovaSeq 6000 | 64970 |
| SRR10488360 | RNA-seq | UIB - sorted uninfected cells from infected well; Canis lupus familiaris; RNA-Seq | Illumina NovaSeq 6000 | 59454 |
| SRR10488361 | RNA-seq | UIC - sorted uninfected cells from infected well; Canis lupus familiaris; RNA-Seq | Illumina NovaSeq 6000 | 49576 |
| SRR10488362 | RNA-seq | UID - sorted uninfected cells from infected well; Canis lupus familiaris; RNA-Seq | Illumina NovaSeq 6000 | 57312 |

| Accession | Data_type | Sample information | Instrument model | Bases count (Mb) |
|---|---|---|---|---|
| SRR10488363 | RNA-seq | UUA - uninfected cells from uninfected well; Canis lupus familiaris; RNA-Seq | Illumina NovaSeq 6000 | 51326 |
| SRR10488364 | RNA-seq | UUB - uninfected cells from uninfected well; Canis lupus familiaris; RNA-Seq | Illumina NovaSeq 6000 | 48019 |
| SRR10488365 | RNA-seq | UUC - uninfected cells from uninfected well; Canis lupus familiaris; RNA-Seq | Illumina NovaSeq 6000 | 52289 |
| SRR10488366 | RNA-seq | UUD - uninfected cells from uninfected well; Canis lupus familiaris; RNA-Seq | Illumina NovaSeq 6000 | 58334 |
| SRR7779680 | RNA-seq | RNA-Seq of canine mammary match normal sample | Illumina HiSeq 2500 | 59693 |
| SRR7779678 | RNA-seq | RNA-Seq of canine mammary match normal sample | Illumina HiSeq 2500 | 113680 |
| SRR7779674 | RNA-seq | RNA-Seq of canine mammary match normal sample | Illumina HiSeq 2500 | 69959 |
| SRR7779671 | RNA-seq | RNA-Seq of canine mammary match normal sample | Illumina HiSeq 2500 | 59369 |
| SRR7779668 | RNA-seq | RNA-Seq of canine mammary match normal sample | Illumina HiSeq 2500 | 65915 |
| SRR7779658 | RNA-seq | RNA-Seq of canine mammary match normal sample | Illumina HiSeq 2500 | 96127 |
| SRR7779652 | RNA-seq | RNA-Seq of canine mammary match normal sample | Illumina HiSeq 2500 | 79805 |
| SRR7779650 | RNA-seq | RNA-Seq of canine mammary match normal sample | Illumina HiSeq 2500 | 83229 |
| SRR7779646 | RNA-seq | RNA-Seq of canine mammary match normal sample | Illumina HiSeq 2500 | 88079 |
| SRR7779642 | RNA-seq | RNA-Seq of canine mammary match normal sample | Illumina HiSeq 2500 | 63686 |
| SRR7779637 | RNA-seq | RNA-Seq of canine mammary match normal sample | Illumina HiSeq 2500 | 71918 |
| SRR7779636 | RNA-seq | RNA-Seq of canine mammary match normal sample | Illumina HiSeq 2500 | 69984 |
| SRR7779632 | RNA-seq | RNA-Seq of canine mammary match normal sample | Illumina HiSeq 2500 | 81060 |
| SRR7779629 | RNA-seq | RNA-Seq of canine mammary match normal sample | Illumina HiSeq 2500 | 78150 |
| SRR7779622 | RNA-seq | RNA-Seq of canine mammary match normal sample | Illumina HiSeq 2500 | 72788 |
| SRR7779617 | RNA-seq | RNA-Seq of canine mammary match normal sample | Illumina HiSeq 2500 | 72344 |
| SRR7779610 | RNA-seq | RNA-Seq of canine mammary match normal sample | Illumina HiSeq 2500 | 57599 |
| SRR7779597 | RNA-seq | RNA-Seq of canine mammary match normal sample | Illumina HiSeq 2500 | 70941 |
| SRR7779577 | RNA-seq | RNA-Seq of canine mammary match normal sample | Illumina HiSeq 2500 | 68303 |
| SRR7779569 | RNA-seq | RNA-Seq of canine mammary match normal sample | Illumina HiSeq 2500 | 57341 |
| SRR7779567 | RNA-seq | RNA-Seq of canine mammary match normal sample | Illumina HiSeq 2500 | 116248 |
| SRR7779564 | RNA-seq | RNA-Seq of canine mammary match normal sample | Illumina HiSeq 2500 | 92521 |
| SRR7779560 | RNA-seq | RNA-Seq of canine mammary match normal sample | Illumina HiSeq 2500 | 63529 |
| SRR7779549 | RNA-seq | RNA-Seq of canine mammary match normal sample | Illumina HiSeq 2500 | 67172 |
| SRR7779544 | RNA-seq | RNA-Seq of canine mammary match normal sample | Illumina HiSeq 2500 | 81885 |
| SRR7779541 | RNA-seq | RNA-Seq of canine mammary match normal sample | Illumina HiSeq 2500 | 67930 |
| SRR7779531 | RNA-seq | RNA-Seq of canine mammary match normal sample | Illumina HiSeq 2500 | 74583 |
| SRR7779525 | RNA-seq | RNA-Seq of canine mammary match normal sample | Illumina HiSeq 2500 | 86119 |
| SRR7779524 | RNA-seq | RNA-Seq of canine mammary match normal sample | Illumina HiSeq 2500 | 78086 |
| SRR7779520 | RNA-seq | RNA-Seq of canine mammary match normal sample | Illumina HiSeq 2500 | 61051 |
| SRR7779514 | RNA-seq | RNA-Seq of canine mammary match normal sample | Illumina HiSeq 2500 | 69656 |
| SRR7779512 | RNA-seq | RNA-Seq of canine mammary match normal sample | Illumina HiSeq 2500 | 62373 |
| SRR7779506 | RNA-seq | RNA-Seq of canine mammary match normal sample | Illumina HiSeq 2500 | 80551 |
| SRR7779496 | RNA-seq | RNA-Seq of canine mammary match normal sample | Illumina HiSeq 2500 | 60064 |
| SRR7779490 | RNA-seq | RNA-Seq of canine mammary match normal sample | Illumina HiSeq 2500 | 78250 |
| SRR7779478 | RNA-seq | RNA-Seq of canine mammary match normal sample | Illumina HiSeq 2500 | 79664 |
| SRR7779476 | RNA-seq | RNA-Seq of canine mammary match normal sample | Illumina HiSeq 2500 | 74245 |
| SRR7779473 | RNA-seq | RNA-Seq of canine mammary match normal sample | Illumina HiSeq 2500 | 69048 |
| SRR7779463 | RNA-seq | RNA-Seq of canine mammary match normal sample | Illumina HiSeq 2500 | 81964 |
| SRR8003175 | RNA-seq | RNA-Seq of canis lupus familiaris: XLMTM adult biceps femoris | Illumina HiSeq 4000 | 34671 |
| SRR8003171 | RNA-seq | RNA-Seq of canis lupus familiaris: XLMTM adult vastus lateralis | Illumina HiSeq 4000 | 176895 |
| SRR8003170 | RNA-seq | RNA-Seq of canis lupus familiaris: WT adult vastus lateralis | Illumina HiSeq 4000 | 198212 |
| SRR8003169 | RNA-seq | RNA-Seq of canis lupus familiaris: XLMTM adult vastus lateralis | Illumina HiSeq 4000 | 177058 |
| SRR8003165 | RNA-seq | RNA-Seq of canis lupus familiaris: WT adult vastus lateralis | Illumina HiSeq 4000 | 197156 |
| ERR2732073 | RNA-seq | Curly Coated Retriever | Illumina HiSeq 3000 | 10024 |
| ERR2732074 | RNA-seq | Curly Coated Retriever | Illumina HiSeq 3000 | 8490 |
| ERR2732075 | RNA-seq | Curly Coated Retriever | Illumina HiSeq 3000 | 10273 |
| ERR2732076 | RNA-seq | Curly Coated Retriever | Illumina HiSeq 3000 | 9397 |
| ERR2732077 | RNA-seq | Curly Coated Retriever | Illumina HiSeq 3000 | 8867 |
| ERR2732079 | RNA-seq | Curly Coated Retriever | Illumina HiSeq 3000 | 8249 |
| ERR2732081 | RNA-seq | Curly Coated Retriever | Illumina HiSeq 3000 | 10910 |

| Accession | Data_type | Sample information | Instrument model | Bases count (Mb) |
|---|---|---|---|---|
| ERR2732082 | RNA-seq | Curly Coated Retriever | Illumina HiSeq 3000 | 11006 |
| ERR2733133 | RNA-seq | Curly Coated Retriever | Illumina HiSeq 3000 | 8683 |
| ERR3274932 | RNA-seq | Labrador Retrievers | Illumina HiSeq 2500 | 11115 |
| ERR3274933 | RNA-seq | Labrador Retrievers | Illumina HiSeq 2500 | 11234 |
| ERR3274937 | RNA-seq | Labrador Retrievers | Illumina HiSeq 2500 | 13977 |
| ERR1948875 | RNA-seq | Transcriptomics of morphologically diverse dog breeds | Illumina HiSeq 4000 | 28688 |
| ERR1948876 | RNA-seq | Transcriptomics of morphologically diverse dog breeds | Illumina HiSeq 4000 | 29069 |
| ERR1948877 | RNA-seq | Transcriptomics of morphologically diverse dog breeds | Illumina HiSeq 4000 | 30978 |
| ERR1948878 | RNA-seq | Transcriptomics of morphologically diverse dog breeds | Illumina HiSeq 4000 | 25749 |
| ERR1948879 | RNA-seq | Transcriptomics of morphologically diverse dog breeds | Illumina HiSeq 4000 | 22676 |
| ERR1948880 | RNA-seq | Transcriptomics of morphologically diverse dog breeds | Illumina HiSeq 4000 | 30154 |
| ERR1948881 | RNA-seq | Transcriptomics of morphologically diverse dog breeds | Illumina HiSeq 4000 | 32341 |
| ERR1948882 | RNA-seq | Transcriptomics of morphologically diverse dog breeds | Illumina HiSeq 4000 | 27088 |
| ERR1948883 | RNA-seq | Transcriptomics of morphologically diverse dog breeds | Illumina HiSeq 4000 | 30758 |

Table S12 Public resource of illumina short-read data of 25 dogs

| Sample ID | Number of paired reads | Reads length (bp) | Sequencing platform | Accession | Genome coverage | Breed |
|---|---|---|---|---|---|---|
| ISR_BOX_1 | 285156478 | 125 | Illumina HiSeq 2500 | ERR2196023 | 28.36X | Boxer |
| ISR_CKCS_1 | 276885105 | 125 | Illumina HiSeq 2500 | ERR2196025 | 27.62X | Cavalier King Charles Spaniel |
| ISR_DH_1 | 364316971 | 150 | Illumina HiSeq 2000 | ERR3047535 | 43.06X | Dachshund |
| ISR_PUG_1 | 344029880 | 150 | Illumina HiSeq 2000 | ERR3047544 | 40.88X | Pug |
| ISR_LEO_1 | 203589257 | 100 | Illumina HiSeq 3000 | SRR7107523 | 16.22X | Leonberger |
| ISR_CS_1 | 222108500 | 150 | Illumina HiSeq 2500 | SRR7107632 | 26.42X | Cocker spaniel |
| ISR_WM_1 | 192335020 | 150 | Illumina HiSeq 2500 | SRR7107634 | 22.96X | Weimaraner |
| ISR_GSD_1 | 194141819 | 100 | Illumina HiSeq 2000 | SRR7107763 | 15.55X | German Shepherd |
| ISR_GSD_2 | 188797150 | 100 | Illumina HiSeq 2000 | SRR7107767 | 14.92X | German Shepherd |
| ISR_GSD_3 | 194166764 | 100 | Illumina HiSeq 2000 | SRR7107769 | 15.53X | German Shepherd |
| ISR_CS_2 | 176178243 | 100 | Illumina HiSeq 2000 | SRR7107863 | 13.71X | English cocker spaniel |
| ISR_SCH_1 | 218290866 | 100 | Illumina HiSeq 2000 | SRR7107866 | 16.77X | Standard Schnauzer |
| ISR_SS_1 | 406968239 | 100 | Illumina HiSeq 2000 | SRR7107883 | 30.84X | English Springer Spaniel |
| ISR_SP_1 | 276357019 | 100 | Illumina HiSeq 2000 | SRR7107900 | 21.71X | Standard Poodle |
| ISR_DBM_1 | 372622195 | 100 | Illumina HiSeq 2000 | SRR7107901 | 28.66X | Doberman |
| ISR_GR_1 | 376930517 | 100 | Illumina HiSeq 2000 | SRR7107926 | 28.91X | Golden retriever |
| ISR_GD_1 | 250616106 | 100 | Illumina HiSeq 2500 | SRR7107941 | 20.21X | Great Dane |
| ISR_BM_1 | 473640347 | 125 | Illumina HiSeq 2500 | SRR7120127 | 47.35X | Bernese Mountain Dog |
| ISR_FCR_1 | 410367234 | 125 | Illumina HiSeq 2500 | SRR7120158 | 40.71X | Flat-coated retriever |
| ISR_RTW_1 | 241991725 | 150 | Illumina HiSeq 2500 | SRR7120208 | 21.81X | Rottweiler |
| ISR_RTW_2 | 502519042 | 125 | Illumina HiSeq 2500 | SRR7120209 | 50.16X | Rottweiler |
| ISR_BOX_2 | 320189252 | 100 | Illumina HiSeq 2000 | SRR8541911 | 23.03X | Boxer |
| ISR_IWH_1 | 370755703 | 100 | Illumina HiSeq 2000 | SRR8541930 | 23.88X | Irish Wolfhound |
| ISR_IWH_2 | 377090701 | 100 | Illumina HiSeq 2000 | SRR8541931 | 24.4X | Irish Wolfhound |
| ISR_CS_3 | 208440846 | 100 | Illumina HiSeq 2500 | SRR8614081 | 15.79X | English cocker spaniel |

Table S13 Summary of dark regions detected in each dog

| Dataset | Dog_id | COV_cutoff | COV length (bp) | CAM length (bp) |
|---|---|---|---|---|
| 10X linked reads | BOX_1 | 4X | 63504052 | 24865150 |
| 10X linked reads | CS_1 | 5X | 62957286 | 22983375 |
| 10X linked reads | FCR_2 | 5X | 61616193 | 23349025 |
| 10X linked reads | SCH_1 | 4X | 63958627 | 22805750 |
| 10X linked reads | BOX_2 | 3X | 65529900 | 25691300 |
| 10X linked reads | FCR_1 | 3X | 60370631 | 26127713 |
| 10X linked reads | GR_1 | 4X | 60319502 | 24283200 |
| 10X linked reads | Mischka | 5X | 43327368 | 21499125 |
| 10X linked reads | BM_1 | 4X | 63360345 | 24476700 |
| 10X linked reads | CKCS_1 | 3X | 62763180 | 25198938 |
| 10X linked reads | CS_2 | 5X | 64299293 | 24154575 |
| 10X linked reads | CS_3 | 5X | 42993327 | 67461900 |
| 10X linked reads | DH_1 | 3X | 60090475 | 25296925 |
| 10X linked reads | DBM_1 | 5X | 51987907 | 27865700 |
| 10X linked reads | GSD_1 | 5X | 42012114 | 47627288 |
| 10X linked reads | GSD_2 | 5X | 51510014 | 36763813 |
| 10X linked reads | GSD_3 | 4X | 60567750 | 23451975 |
| 10X linked reads | GD_1 | 4X | 60089132 | 25074013 |
| 10X linked reads | IWH_1 | 3X | 60890152 | 26493375 |
| 10X linked reads | IWH_2 | 4X | 65090150 | 25018600 |
| 10X linked reads | LEO_1 | 3X | 61447227 | 25888450 |
| 10X linked reads | PUG_1 | 3X | 64122768 | 26481925 |
| 10X linked reads | RTW_1 | 5X | 59165212 | 51601738 |
| 10X linked reads | RTW_2 | 5X | 63844831 | 23224171 |
| 10X linked reads | SS_1 | 5X | 54582377 | 37377875 |
| 10X linked reads | SBT_1 | 4X | 65644202 | 23527750 |
| 10X linked reads | SP_1 | 5X | 56273459 | 33689238 |
| 10X linked reads | WM_1 | 5X | 58933527 | 23413500 |
| Illumina short reads | ISR_BOX_1 | 4X | 60154879 | 64639179 |
| Illumina short reads | ISR_CKCS_1 | 4X | 61432287 | 65637933 |
| Illumina short reads | ISR_DH_1 | 5X | 51614454 | 64646045 |
| Illumina short reads | ISR_PUG_1 | 5X | 57190960 | 64634602 |
| Illumina short reads | ISR_LEO_1 | 2X | 67061803 | 81226347 |
| Illumina short reads | ISR_CS_1 | 4X | 60789790 | 64113222 |
| Illumina short reads | ISR_WM_1 | 3X | 63525773 | 65682781 |
| Illumina short reads | ISR_GSD_1 | 2X | 60893621 | 88900970 |
| Illumina short reads | ISR_GSD_2 | 2X | 60409830 | 88095117 |
| Illumina short reads | ISR_GSD_3 | 2X | 60120901 | 89075442 |
| Illumina short reads | ISR_CS_2 | 3X | 65438931 | 70224697 |
| Illumina short reads | ISR_SCH_1 | 4X | 68118028 | 67762656 |

| Dataset | Dog_id | COV_cutoff | COV length (bp) | CAM length (bp) |
|---|---|---|---|---|
| Illumina short reads | ISR_SS_1 | 5X | 57009176 | 83826854 |
| Illumina short reads | ISR_SP_1 | 4X | 62904016 | 75147087 |
| Illumina short reads | ISR_DBM_1 | 4X | 62243607 | 82862506 |
| Illumina short reads | ISR_GR_1 | 5X | 55578516 | 75584105 |
| Illumina short reads | ISR_GD_1 | 5X | 64102203 | 72796234 |
| Illumina short reads | ISR_BM_1 | 5X | 50687689 | 70471503 |
| Illumina short reads | ISR_FCR_1 | 5X | 55634513 | 68047817 |
| Illumina short reads | ISR_RTW_1 | 2X | 66203017 | 70920792 |
| Illumina short reads | ISR_RTW_2 | 5X | 51014475 | 69883500 |
| Illumina short reads | ISR_BOX_2 | 3X | 60418596 | 99966997 |
| Illumina short reads | ISR_IWH_1 | 5X | 64218973 | 93560359 |
| Illumina short reads | ISR_IWH_2 | 5X | 63408979 | 94060469 |
| Illumina short reads | ISR_CS_3 | 3X | 60235778 | 71036808 |
| PacBio long reads | Mischka | 5X | 6442451 | 1043275 |

Table S14. Tissues samples and genotyping results for expression analysis

| Tissue | ID | Breed | Sex[2] | Age[3] | RIN | Genotype[1] | | | | | |
|---|---|---|---|---|---|---|---|---|---|---|---|
| | | | | | | *POLI* | *MANEA* | *RAB32* | *CYP1A2* | *PPHLN1* | *OTOA* |
| Liver | 1L | Danish Swedish Farmdog | F | 12 | 7.3 | na | het | del | 4 | 2 | 2 |
| | 10L | German Shepherd | F | 12 | 8.1 | na | het | del | 6 | 2 | 2 |
| | 11 L | Irish Setter | M | 6 | 7.7 | na | del | wt | 3 | 2 | 2 |
| | 12 L | Golden Retriever | F | 11 | 7.7 | na | del | wt | 4 | 4 | 2 |
| | 13 L | Poodle Medium | F | 13 | 7.2 | na | del | wt | 2 | 2 | 2 |
| | 17 L | Hovawart | F | 7 | 8.6 | na | het | wt | 4 | 3 | 2 |
| | 18 L | German Shepherd | F | 12 | 5.9 | na | het | wt | 3 | 2 | 3 |
| | 2 L | Mixed Breed | na | na | 7.2 | na | het | na | 5 | 2 | 4 |
| | 20 L | Beagle | M | na | 3.6 | na | del | wt | 3 | 3 | 2 |
| | 22 L | German Shepherd | F | 5 | 7.6 | na | het | wt | 4 | 2 | 2 |
| | 23 L | Swedish Elkhound | F | 9 | 6.5 | na | het | wt | 3 | 2 | 2 |
| | 24 L | German Shepherd | F | 1 | 8 | na | het | wt | 3 | 3 | 2 |
| | 25 L | German Shepherd | M | 1 | 8.3 | na | het | wt | 6 | 3 | 2 |
| | 3 L | Howawart | F | 11 | 8 | na | het | wt | 4 | 3 | 2 |
| | 4 L | Howawart | M | 11 | 6.8 | na | het | wt | 4 | 2 | 2 |
| | 5 L | German Shepherd | M | 12 | 7.6 | na | del | del | 3 | 2 | 2 |
| | 6 L | Irish Setter | M | na | 7.9 | na | del | wt | 4 | 2 | 2 |
| | 7 L | Greyhound | F | 12.5 | 7.6 | na | het | wt | 3 | 2 | 2 |
| | 8 L | Border Collie | F | 13 | 8.1 | na | het | wt | 3 | 3 | 2 |
| | 9 L | Howawart | F | 9 | 6 | na | het | wt | 5 | 2 | 2 |
| Spleen | 1 S | Cocker Spaniel | M | 9 | 8.1 | na | het | wt | 4 | 2 | 2 |
| | 10S | German Shepherd | M | 12 | 7.3 | na | del | del | 3 | 3 | 2 |
| | 11 S | Howawart | F | 11 | 8.7 | na | het | wt | 4 | 3 | 2 |
| | 12 S | German Shepherd | F | 12 | 6 | na | het | wt | 3 | 2 | 3 |
| | 14 S | Beagle | M | na | 6.2 | na | del | wt | 3 | 3 | 2 |
| | 15 S | German Shepherd | F | 5 | 7.5 | na | het | wt | 4 | 2 | 2 |
| | 16 S | German Shepherd | F | 1 | 7.7 | na | het | wt | 3 | 3 | 2 |
| | 17 S | German Shepherd | M | 1 | 7.4 | na | het | wt | 6 | 3 | 2 |
| | 2 S | Irish Setter | M | na | 8.8 | na | del | wt | 4 | 2 | 2 |
| | 3 S | Dachshund | F | 15 | 9.3 | na | het | wt | 4 | 2 | 2 |
| | 4 S | Danish Swedish Farmdog | F | 12 | 9.2 | na | het | del | 4 | 2 | 2 |
| | 5 S | Border Collie | F | 13 | 7.8 | na | het | wt | 3 | 2 | 2 |
| | 6 S | German Shepherd | F | 2 | 8.5 | na | het | wt | 5 | 3 | 2 |
| | 7 S | Poodle Medium | F | 13 | 6.6 | na | del | wt | 2 | 2 | 2 |

| Tissue | ID | Breed | Sex[2] | Age[3] | RIN | Genotype[1] | | | | | |
|---|---|---|---|---|---|---|---|---|---|---|---|
| | | | | | | *POLI* | *MANEA* | *RAB32* | *CYP1A2* | *PPHLN1* | *OTOA* |
| | 8 S | Golden Retriever | F | 11 | 7.7 | na | del | wt | 4 | 4 | 2 |
| Left Ventricle Wall | c10 | Cavalier King Charles Spaniel | M | 9 | 8.4 | del | na | na | na | na | na |
| | c11 | Cavalier King Charles Spaniel | F | 9 | 8.2 | del | na | na | na | na | na |
| | c9 | Cavalier King Charles Spaniel | F | 9 | 8.4 | del | na | na | na | na | na |
| | c2 | Cavalier King Charles Spaniel | M | 10 | 8.8 | het | na | na | na | na | na |
| | c7 | Cavalier King Charles Spaniel | F | 11 | 8.9 | het | na | na | na | na | na |
| | c8 | Cavalier King Charles Spaniel | F | 12 | 8.4 | het | na | na | na | na | na |
| | d3 | Dobermann | F | 5 | 8.8 | wt | na | na | na | na | na |
| | d8 | Dobermann | F | 9 | 8.2 | wt | na | na | na | na | na |
| | d9 | Dobermann | M | 6 | 8.1 | wt | na | na | na | na | na |

1Genotype: wt, wild type; het, heterozygote; del, deletion; CNV as binned whole numbers. 2 Sex: F, female; M, male. 3Age in years. na, not available
